# Small molecule activation of metabolic enzyme pyruvate kinase muscle isozyme 2, PKM2, provides photoreceptor neuroprotection

**DOI:** 10.1101/797118

**Authors:** Thomas J. Wubben, Mercy Pawar, Eric Weh, Andrew Smith, Peter Sajjakulnukit, Li Zhang, Lipeng Dai, Heather Hager, Manjunath P. Pai, Costas A. Lyssiotis, Cagri G. Besirli

## Abstract

Photoreceptor cell death is the ultimate cause of vision loss in many retinal disorders, and there is an unmet need for neuroprotective modalities to improve photoreceptor survival. Similar to cancer cells, photoreceptors maintain pyruvate kinase muscle isoform 2 (PKM2) expression, which is a critical regulator in aerobic glycolysis. Unlike PKM1, which has constitutively high catalytic activity, PKM2 is under complex regulation. Recently, we demonstrated that genetically reprogramming photoreceptor metabolism via PKM2-to-PKM1 substitution is a promising neuroprotective strategy. Here, we explored the neuroprotective effects of pharmacologically activating PKM2 via ML-265, a small molecule activator of PKM2, during acute outer retinal stress. We found that ML-265 increased PKM2 activity in 661W cells and *in vivo* in rat eyes without affecting the expression of genes involved in glucose metabolism. ML-265 treatment did, however, alter metabolic intermediates of glucose metabolism and those necessary for biosynthesis in cultured cells. Long-term exposure to ML-265 did not result in decreased photoreceptor function or survival under baseline conditions. Notably, though, ML-265-treatment did reduce entrance into the apoptotic cascade in *in vitro* and *in vivo* models of outer retinal stress. These data suggest that reprogramming metabolism via activation of PKM2 is a novel, and promising, therapeutic strategy for photoreceptor neuroprotection.

## Introduction

Photoreceptor cell death is the ultimate cause of vision loss in many retinal disorders, including retinal detachment, retinal dystrophies, and non-exudative age-related macular degeneration. Unfortunately, no successful treatment options currently exist to prevent this cell loss. Therefore, there is an urgent unmet need for neuroprotective modalities to improve photoreceptor survival. Interestingly, mutations affecting enzymes critical for energy and purine metabolism, common to almost all cells in the body, have been linked to isolated retinal degenerations.^1^ Considering photoreceptors have little reserve capacity to generate adenosine triphosphate (ATP), and as a result, are susceptible to small changes in energy homeostasis, disruption of nutrient availability and metabolic regulation may be a unifying mechanism in photoreceptor cell death.^1–3^ Hence, an improved understanding of the metabolic signals that regulate photoreceptor survival during outer retinal stress may identify novel targets for developing neuroprotective therapies for a plethora of retinal diseases.

Similar to cells with high metabolic demands, such as cancer cells, photoreceptors are capable of aerobic glycolysis.^4–6^ Aerobic glycolysis is the conversion of glucose to lactate despite the presence of oxygen. In cancer, it is thought that this process and its regulatory enzymes provide the tumor cells with the flexibility to adapt their metabolism to a state conducive to growth and proliferation or energy production during periods of nutrient deprivation.^7,8^ Recent studies have suggested that aerobic glycolysis and its essential regulatory enzymes, hexokinase 2 (HK2), pyruvate kinase muscle isoform 2 (PKM2), and lactate dehydrogenase A (LDHA), support photoreceptor outer segment biogenesis as well as fuel Müller cells and retinal pigment epithelium (RPE) so that sufficient glucose can nourish the outer retina.^5,9,10^ Furthermore, aerobic glycolysis and its key enzymes have been shown to be important for the function and survival of rod and cone photoreceptors under baseline conditions and confer a survival advantage to photoreceptors during periods of outer retinal stress.^4,11–14^

We recently reported that a key regulator of aerobic glycolysis, PKM2, plays a critical function in photoreceptor survival during acute outer retinal stress.^15^ PKM2 converts phosphoenolpyruvate (PEP) and adenosine diphosphate (ADP) to pyruvate and ATP in the final rate-limiting step of glycolysis. Photoreceptors maintain expression of PKM2 after differentiation, unlike other neurons, which exclusively express the constitutively active PKM1 isoform following terminal differentation.^5,13,15–17^ PKM2 activity is tightly regulated whereas PKM1 is constitutively active.^8^ As a tetramer, PKM2 displays high catalytic activity and is associated with increased ATP synthesis and catabolic metabolism. This quaternary state is favored by nutrient deprivation and certain allosteric regulators, such as serine and fructose-1,6-bisphosphate.^8^ PKM2 can also exist in a non-tetrameric form, which has a low catalytic activity and is associated with anabolic metabolism and the shuttling of metabolic intermediates to biosynthetic pathways.^8,18,19^ This catalytic state is supported by nutrient availability, growth factors, and post-translational modifications, such as phosphorylation.^8^

A previous study in our lab explored the role of PKM2 in photoreceptors using a rod photoreceptor-specific, *Pkm2* conditional knockout mouse model (Rho-Cre:*Pkm2*^*f/f*^). PKM2 localizes primarily to the outer retina while PKM1 expression is mostly confined to the inner retina.^12,15,20^ The loss of *Pkm2* in photoreceptors led to the compensatory expression of *Pkm1* and upregulation of genes involved in glucose metabolism. When these animals were subjected to acute outer retinal stress produced by experimental retinal detachment, they displayed decreased photoreceptor apoptosis and improved survival. We also determined that the phosphorylation state of PKM2 in wild-type rodent retina decreases after experimental retinal detachment implying an increase in catabolic activity. Thus, the genetic PKM2-to-PKM1 isoform shift observed in the photoreceptors of Rho-Cre:*Pkm2*^*f/f*^ mice mimics the activation of PKM2 after nutrient deprivation by substituting constitutively active PKM1 to circumvent acute apoptotic stress. These data suggest that increasing PKM2 catalytic activity using a small-molecule activator may show similar photoreceptor neuroprotective effects. ML-265 (also known as TEPP-46) has been extensively utilized in the literature as a potent activator of PKM2 and was originally developed as a potential cancer therapy.^21–24^ ML-265 was shown to increase PK activity in cells expressing mainly PKM2 to levels comparable to cells expressing only the PKM1 isoform, demonstrating that ML-265-mediated activation of PKM2 mimics a PKM2-to-PKM1 isoform shift similar to that seen in our mouse model.^22^ Therefore, ML-265 represents an ideal lead molecule to explore the neuroprotective effects of pharmacologically activating PKM2 in photoreceptors to reprogram photoreceptor metabolism.

In the present study, we explored the neuroprotective effects of pharmacologically reprogramming photoreceptor metabolism by altering PKM2 function with ML-265 during outer retinal apoptotic stress. We found that ML-265 can increase PK activity in the cellular context utilizing 661W cone-like cells *in vitro* as well as *in vivo* via intravitreal injections into rat eyes.^25,26^ Interestingly, increasing total PK activity did not alter the expression of genes involved in glucose metabolism but did affect the intracellular concentrations of metabolites involved in glycolysis and biosynthesis. Despite these metabolic effects, no significant differences in photoreceptor function or survival were observed over time under baseline conditions between ML-265 and vehicle-treated rat eyes. However, in both *in vitro* and *in vivo* models of outer retinal apoptotic stress, ML-265-mediated PKM2 activation reduced entrance into the apoptotic cascade.

## Results

### ML-265 increases pyruvate kinase activity *in vitro* and *in vivo*

While ML-265 has been extensively utilized in the literature as a potent activator of PKM2, its ability to activate PKM2 in the eye has never been examined.^21–24^ PKM2 is expressed by photoreceptors in the retina while PKM1 expression is confined to the inner retina, and ML-265 shows a high degree of selectivity for PKM2 over other isoforms, PKM1, PKR, and PKL. ^5,10,12,13,15,17,20–22^ Therefore, ML-265 represents an ideal lead molecule to explore the neuroprotective effects of pharmacologically activating PKM2 in photoreceptors. In line with previous reports, ML-265 displayed potent activation of recombinant human PKM2 with a maximum activation of 249 ± 14% (best-fit value ± std. error) and a half-maximum activating concentration (AC_50_) of 108 ± 20 nM (Fig. 1a).^21,22^ To begin to investigate the ability of ML-265 to activate PK in photoreceptors, the 661W cell line was utilized. 661W cells are immortalized photoreceptors that express cone markers.^25,27^ Similar to the results observed with the recombinant enzyme, ML-265 increased PK activity *in vitro* (maximum activation=515 ± 12% and AC_50_=19 ± 2 nM) (Fig. 1b). This *in vitro* data suggests the small molecule activator is able to cross the cell membrane and enhance PK activity. With this understanding, intravitreal injections of 2 μL of increasing concentrations of ML-265 or equal volume of vehicle (dimethyl sulfoxide, DMSO) were performed in rats with the retinas harvested four hours after injection and lysate assayed for PK activity (Fig. 1c). Intravitreal injection of ML-265 was able to activate PK *in vivo* up to 170 ± 26% with an AC_50_=59 ± 38 μM. Considering both the specificity of ML-265 for PKM2 and the fact that PKM2 expression is confined to the outer retina, ML-265 is most likely able to traverse the retina and the cell membranes of photoreceptors to activate PKM2.^5,10,15,17,22^

**Figure 1.**
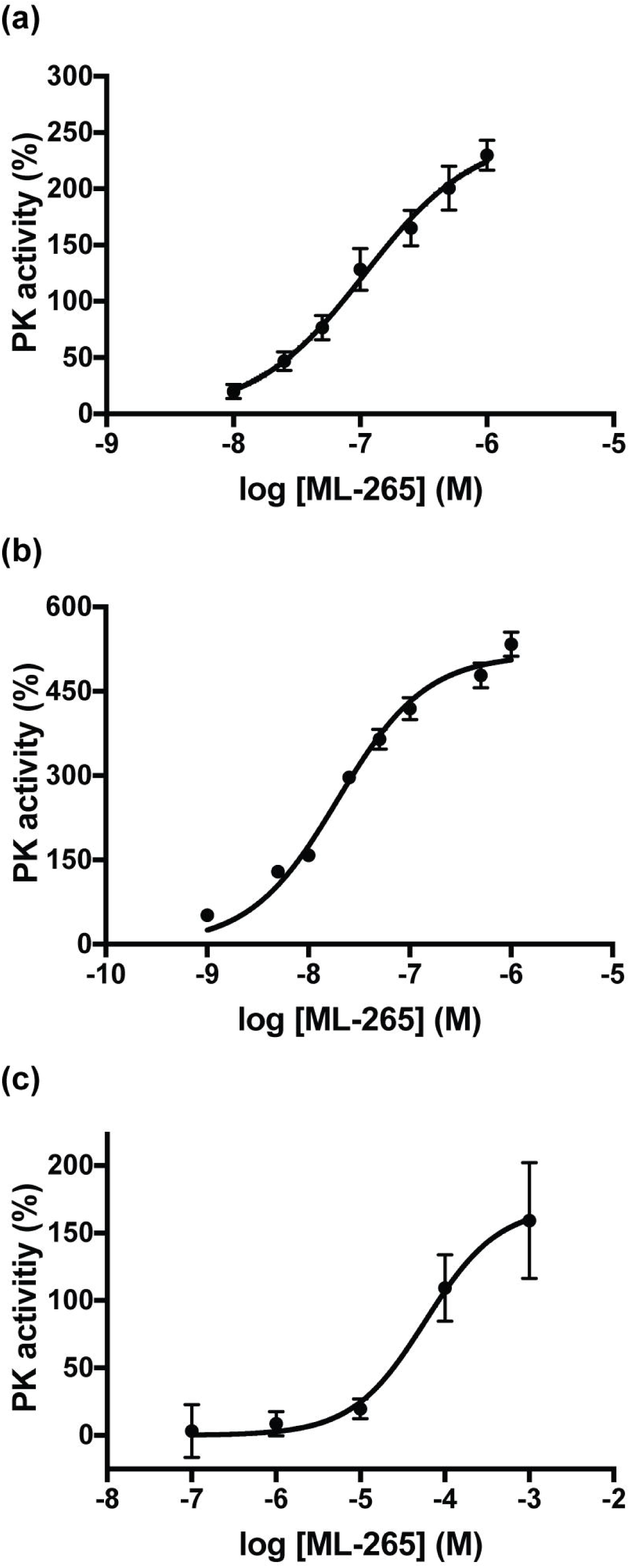
Pharmacologic activation of PKM2 using ML-265 increases PK activity *in vitro* and *in vivo*. Pyruvate kinase (PK) activity in the presence of increasing ML-265 concentrations with either: (a) human recombinant PKM2, (b) 661W cells (n=3 replicates per ML-265 concentration), or (c) rat retinas after intravitreal injections (n=3 animals per ML-265 concentration). Data were normalized to DMSO treated PK activity in recombinant, *in vitro*, and *in vivo* experiments. Mean ± SEM.

### Pharmacokinetic analysis of ML-265 intravitreal injection

Understanding the intraocular pharmacokinetics of an intravitreal injection of ML-265 is essential to future therapeutic strategies for retinal diseases as well as to be able to extend experimentation into the *in vivo* realm. To this end, we determined the intraocular pharmacokinetic profile of intravitreally administered ML-265 in rabbits, which are the most commonly used animals in intravitreal pharmacokinetic studies with good correlation to the human eye.^28^ Single, intravitreal injections (50 μL) of two different doses of ML-265 were performed in rabbits. Assuming a vitreous volume of 1.15 mL, the final concentrations in the rabbit vitreous were approximately 100 μM and 1000 μM, respectively.^28^ The aqueous humor was sampled at multiple time points after the single intravitreal injection. The drug concentration versus time data was best fit by a two-phase profile as observed in the biphasic decline of the predicted elimination profile (Fig. 2a). The first phase corresponds with distribution, and the latter phase corresponds with elimination. The near parallel concentration-time profiles suggest the distribution and elimination processes are not influenced by dose. The distribution phase has a half-life (*t*_*1/2*_) of 8.3 ± 0.5 hours. The elimination phase has a *t*_*1/2*_=141.6 ± 35 hours. Hence, ML-265 may be present in the eye and provide PK activation for over one week. In accordance with this pharmacokinetic profile, after 15 days, a single intravitreal injection of ML-265 in rats continued to produce a median PK activation in the retina of 38% above that of the DMSO treated retinas (Fig. 2b).

**Figure 2.**
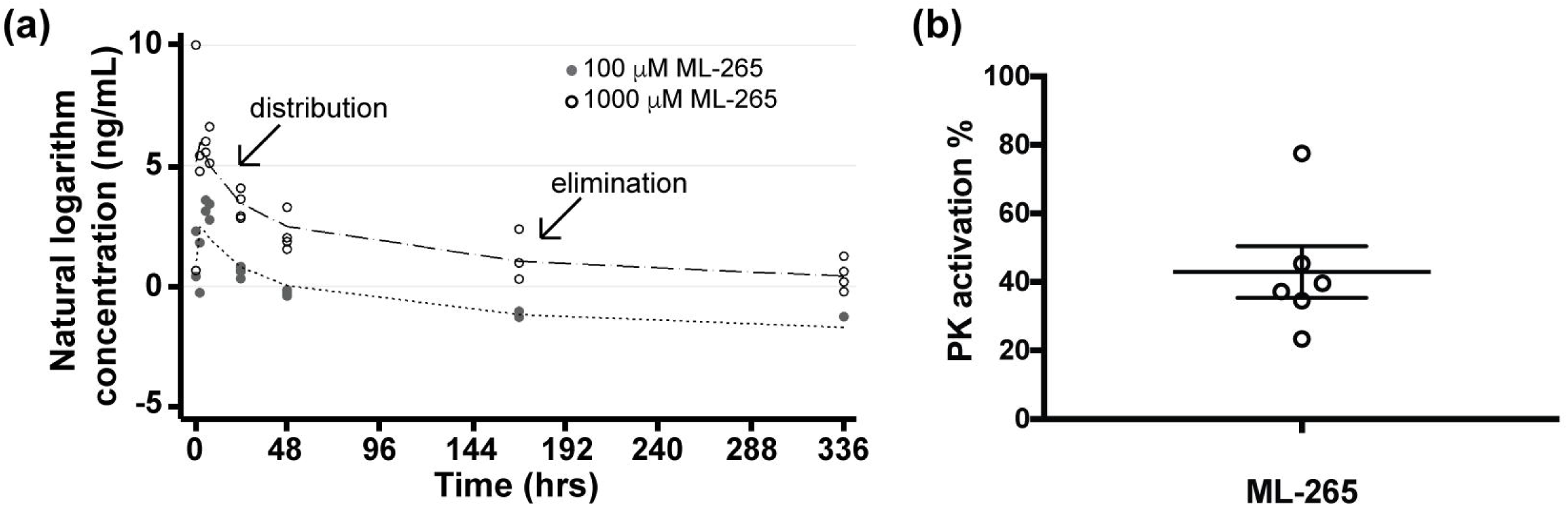
ML-265 has a long half-life and efficacy following intravitreal injection. (a) ML-265 concentration (ng/mL) in rabbit aqueous humor over time after single intravitreal injections of ML-265 (50 μL of 2.3 mM and 23 mM), which provided final vitreous concentrations of 100 μM and 1000 μM, respectively. Dashed lines signify predicted profiles. The distribution and elimination phases are noted on the plot. n=4 eyes per ML-265 concentration. (b) PK activation in harvested rat retinas 15 days after a single intravitreal injection of 2 μL of 7.5 mM ML-265, which equates to 1 mM ML-265 final vitreous concentration, as compared to DMSO treated retinas. Mean ± SEM; n=6 eyes per treatment.

### ML-265 toxicity analysis

ML-265 has been shown to be systemically well-tolerated in mice with over 7 weeks of continuous drug exposure.^21^ However, conditional knockout of *Pkm2* in photoreceptors (Rho-Cre:*Pkm2*^*f/f*^) resulted in upregulation of the constitutively active PKM1 isoform and demonstrated mild degenerative changes in the outer retina and its function over time.^12,15^ ML-265-mediated PKM2 activation mimics a PKM2-to-PKM1 isoform shift, which could cause similar degenerative changes in the retina.^22^ Therefore, a repeat-dose tolerability study was undertaken to determine if prolonged pharmacologic activation of PKM2 would result in anatomic and/or functional changes in the retina. The pharmacokinetic profile of ML-265 suggested that this small-molecule has an intraocular half-life of approximately 6 days (Fig. 2a). As such, weekly intravitreal injections of either a saturating concentration of ML-265 (Fig. 1c) or vehicle (DMSO) were performed in rats for a total of 6 weeks. Optical coherence tomography (OCT) was utilized to measure retinal thickness *in vivo* at baseline immediately prior to the first intravitreal injection and again, one week after the sixth and final injection (Fig. 3a). No statistically significant difference in outer retinal thickness (outer plexiform layer to retinal pigment epithelium inner surface) or outer segment equivalent length (OSEL, inner segment/outer segment junction to retinal pigment epithelium inner surface) was observed after the six weekly, intravitreal injections in the ML-265 or DMSO treated eyes. Nor was a significant difference in outer retinal thickness or OSEL observed between the treatment groups (Fig. 3b and c). Following the final *in vivo* OCT acquisition, the eyes were enucleated for *ex vivo* histological analysis. In accordance with the *in vivo* observations, *ex vivo* analyses (Fig. 3d-f) did not show any statistically significant differences in the number of nuclei contained in the outer nuclear layer (ONL) or total area of the ONL between the ML-265 and DMSO treated eyes after the six, weekly, intravitreal injections.

**Figure 3.**
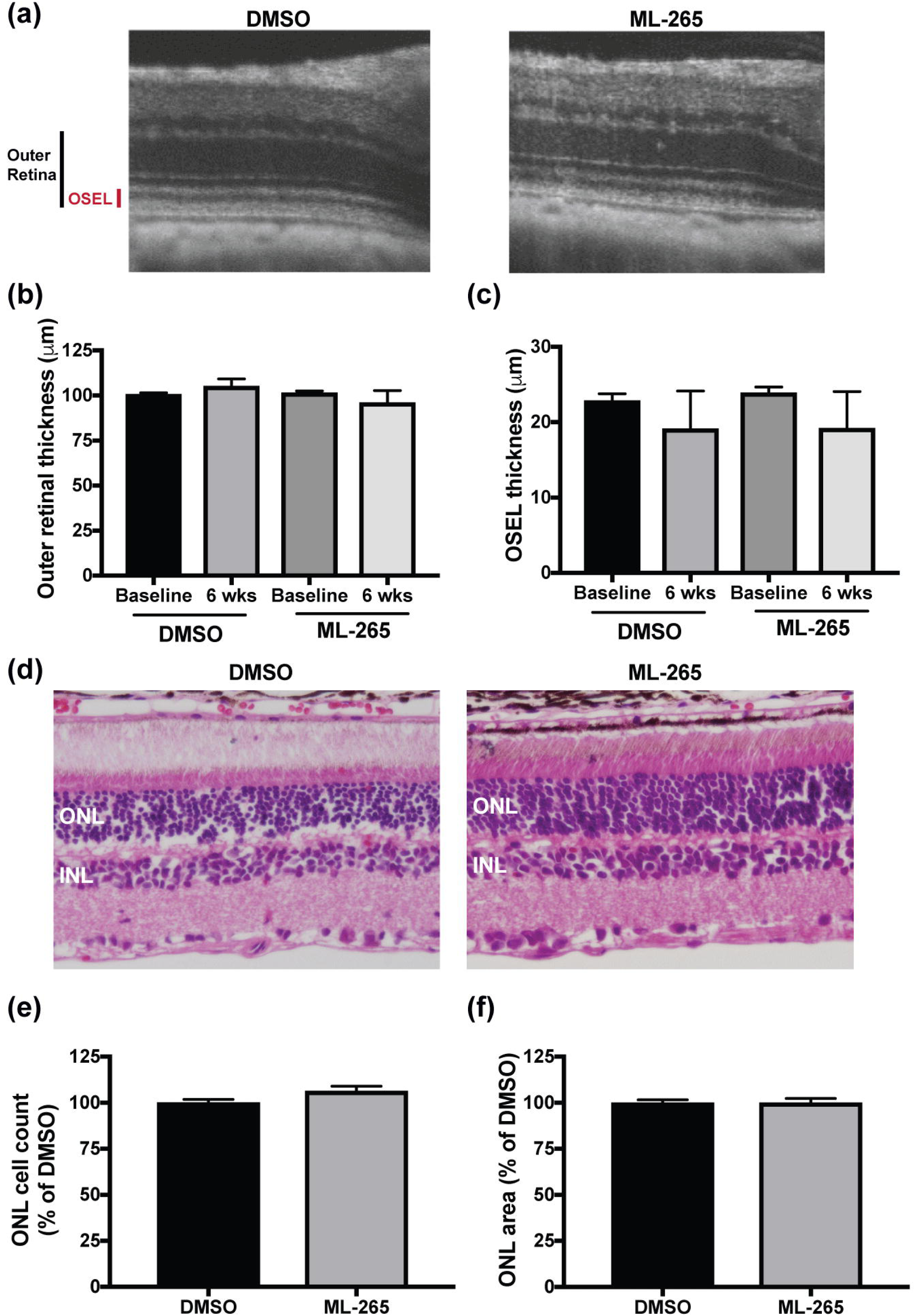
Photoreceptor survival is not affected by long-term ML-265 treatment *in vivo*. (a) Representative OCT images from DMSO and ML-265 treated eyes after 6 weekly intravitreal injections. OSEL, outer segment equivalent length. (b) Outer retinal thickness as determined by OCT in DMSO and ML-265 treated eyes. Measurements were determined immediately prior to starting 6 weekly intravitreal injections (baseline) and again, one week after the last intravitreal injection (6 weeks). (c) OSEL as determined by OCT in DMSO and ML-265 treated eyes. n ≥ 4 eyes per treatment per time point. (d) Representative hematoxylin and eosin stained retinal sections from DMSO and ML-265 treated eyes after 6 weekly intravitreal injections. ONL, outer nuclear layer; INL, inner nuclear layer. (e) ONL cell counts as a percent of the vehicle-treated (DMSO) retinas. Retinas were harvested one week after the last of 6 weekly intravitreal injections with either DMSO or ML-265. (f) ONL area as a percent of the vehicle-treated (DMSO) retinas. n=8 eyes per treatment. Mean ± SEM; *, p<0.05; **, p<0.01; ***, p<0.005.

At the same time, electroretinography (ERG) was utilized to examine if prolonged, pharmacologic activation of PKM2 by ML-265 altered retinal function despite the lack of observed anatomical changes. Interestingly, retinal function, as evaluated by scotopic electroretinography, showed a statistically significant decline in the scotopic a- and b-wave amplitudes after the six weekly, intravitreal injections in both the ML-265 and DMSO treated eyes (Fig. 4a,b, and d). A similar decline was observed in the photopic b-wave in both treatment groups (Fig. 4c and e). However, significant differences were not detected when comparing the baseline electroretinography amplitudes or the 6-week amplitudes between the treatment groups. These data suggest that high concentrations of DMSO (vehicle), and not ML-265, attenuate retinal function.

**Figure 4.**
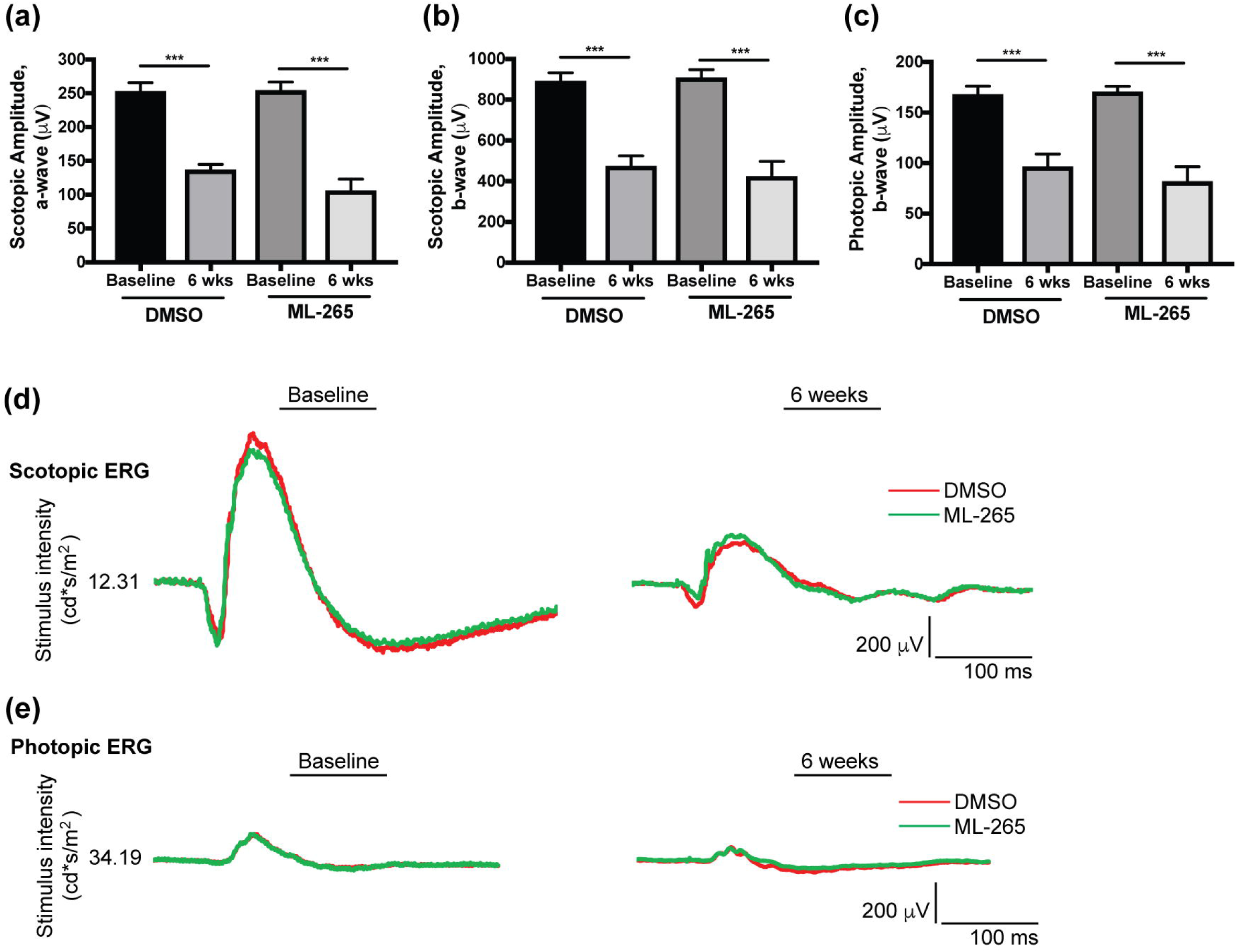
Long-term ML-265 treatment *in vivo* does not impact retinal function. (a) Electroretinography (ERG) scotopic a-wave amplitudes in DMSO and ML-265 treated eyes. Measurements were obtained immediately prior to starting 6 weekly intravitreal injections (baseline) and again, one week after the last intravitreal injection (6 weeks). (b) ERG scotopic b-wave amplitudes in DMSO and ML-265 treated eyes. A flash intensity of 12.31 cd*s/m^2^ was utilized for scotopic amplitudes. (c) ERG photopic b-wave amplitudes in DMSO and ML-265 treated eyes. A flash intensity of 34.19 cd*s/m^2^ was utilized. n ≥ 7 eyes per treatment per time point. Mean ± SEM; *, p<0.05; **, p<0.01; ***, p<0.005. (d) Representative scotopic ERG tracings for DMSO and ML-265 treated eyes at baseline and after the 6 weekly, intravitreal injections. (e) Representative photopic ERG tracings for DMSO and ML-265 treated eyes at baseline and after the 6 weekly, intravitreal injections.

### ML-265 treatment does not alter the expression of genes involved in glucose metabolism

As stated previously, PKM2 is subject to complex regulation, which allows the cell to adapt its metabolic state to different physiologic conditions.^8^ PKM1, which is not under extensive regulation, instead facilitates glycolytic flux, ATP synthesis, and catabolic metabolism.^8,21^ ML-265 activation of PKM2 has been shown to mimic the constitutively active isoform, PKM1.^22^ Therefore, the enhanced activity observed in Figure 1 may induce similar gene expression changes to that previously reported in PKM1-expressing photoreceptors, namely increased expression of genes involved in glucose metabolism.^15^ To determine if ML-265 alters the expression of genes involved in glucose metabolism *in vitro* and *in vivo*, we conducted quantitative reverse transcription PCR using the Glucose Metabolism RT^2^ Profiler™ PCR Array (Qiagen) for either mouse (661W cells) or rat transcripts. This array profiles 84 genes involved in glycolysis, gluconeogenesis, the tricarboxylic acid (TCA) cycle, the pentose phosphate pathway, glycogen synthesis, glycogen degradation, and regulation of glucose and glycogen metabolism. Interestingly, using a fold change cut-off of 2, the normalized gene expression patterns of DMSO and ML-265 treated 661W cells did not show any significant differences after three days of treatment (Fig. 5a). Similarly, rat retinas treated with either an intravitreal injection of ML-265 or equal volume DMSO did not show any significant differences in the gene expression patterns after 3 or 7 days (Fig. 5b and c).

**Figure 5.**
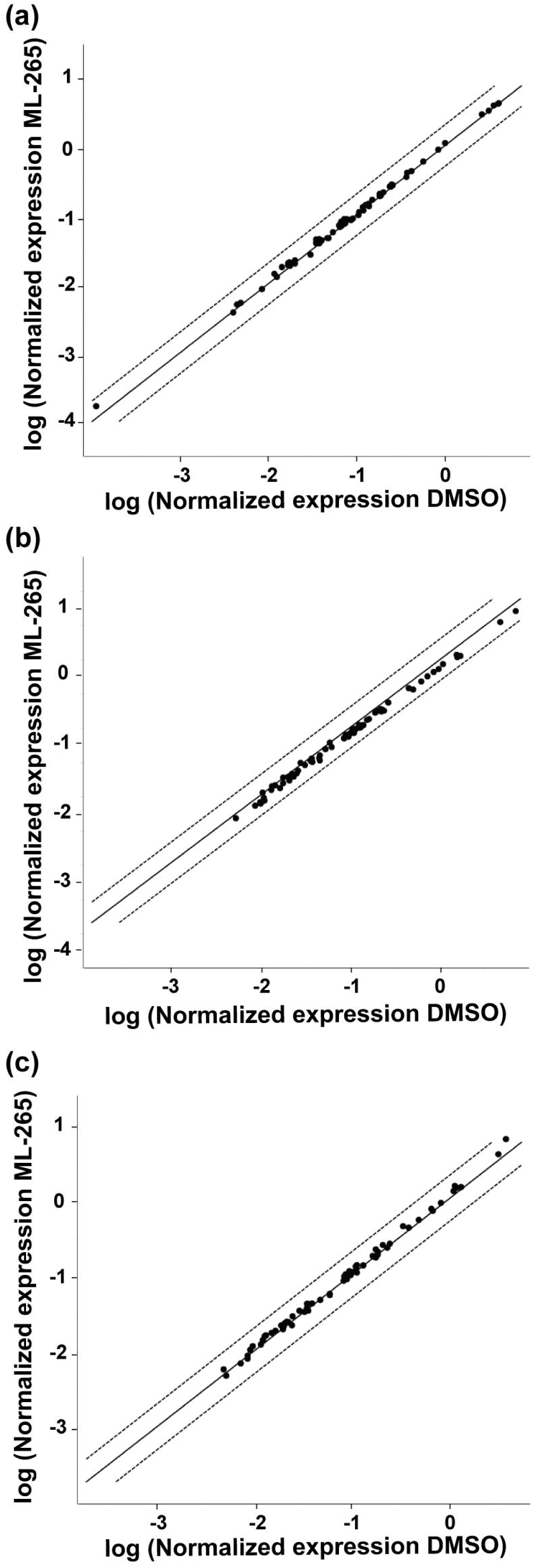
Expression of genes involved in glucose metabolism is unchanged after ML-265 treatment. Scatter plots comparing the normalized expression of genes involved in glucose metabolism between DMSO and ML-265 treated (a) 661W cells after 3 days and rat retinas after (b) 3 and (c) 7 days. The central solid line indicates unchanged gene expression. The dashed outer lines represent the fold change cut-off, which was set to 2. Black solid dots represent those genes that did not surpass the fold change cut-off. n=3 cell replicates or 3-6 animals per group.

### ML-265 treatment alters metabolic profile of cultured photoreceptor-like cells

Since no significant differences in the gene expression patterns were identified after *in vitro* or *in vivo* ML-265 treatment (Fig. 5), we next sought to investigate if ML-265 alters the metabolic profile of cells. Considering ML-265 promotes high PKM2 activity (Fig. 1) and has been shown to mimic the regulatory properties of the constitutively active isoform PKM1, we hypothesized that ML-265 treatment of cultured 661W cells would alter glucose metabolism and the intracellular concentrations of intermediates required for biosynthesis that are derived from glycolysis.^21^ LC-MS/MS was utilized to target 224 metabolites in central glucose, amino acid, nucleotide, and lipid metabolism (Table S1, Supplementary Information).^29,30^ ML-265 treatment induced a decrease in the intracellular concentrations of dihydroxyacetone phosphate (DHAP), glyceraldehyde 3-phosphate (G3P), 2-phosphoglycerate (2-PG), and PEP, which are all glycolytic intermediates proximal to the PKM2-catalyzed reaction (Fig. 6a-c). The ratio of intracellular PEP to pyruvate was statistically significantly decreased in the ML-265 treated cells (Fig. 6d), which further demonstrates the ability of ML-265 to activate PKM2 as PEP is a substrate of this enzyme and pyruvate a product. Consistent with previous studies, a decrease in the intracellular concentrations of ribose 5-phosphate (R5P) and serine (Ser) were also observed (Fig. 6b and c).^21^ R5P is involved in the biosynthesis of nucleotides and is a key intermediate in the pentose phosphate pathway, which is linked to glycolysis upstream of PKM2. Similarly, serine can be synthesized *de novo* from glycolytic intermediates, and is involved in the synthesis of other amino acids and phospholipids, and donates carbon to the folate cycle.^31^ Therefore, these data indicate that ML-265-mediated activation of PKM2 can alter glucose metabolism to produce a more catabolic state. Notably, similar alterations in metabolite pools were not evident in the retinas with PKM1-expressing rod photoreceptors (Rho-Cre:*Pkm2*^*f/f*^) (Fig. 6e). In these retinas, relative to control retinas where both rod *Pkm2* alleles are present (Rho-Cre:*Pkm2*^+*/*+^), there was a statistically significant increase in 2-PG and a trend towards increased PEP, consistent with previous metabolic flux studies.^12^

**Figure 6.**
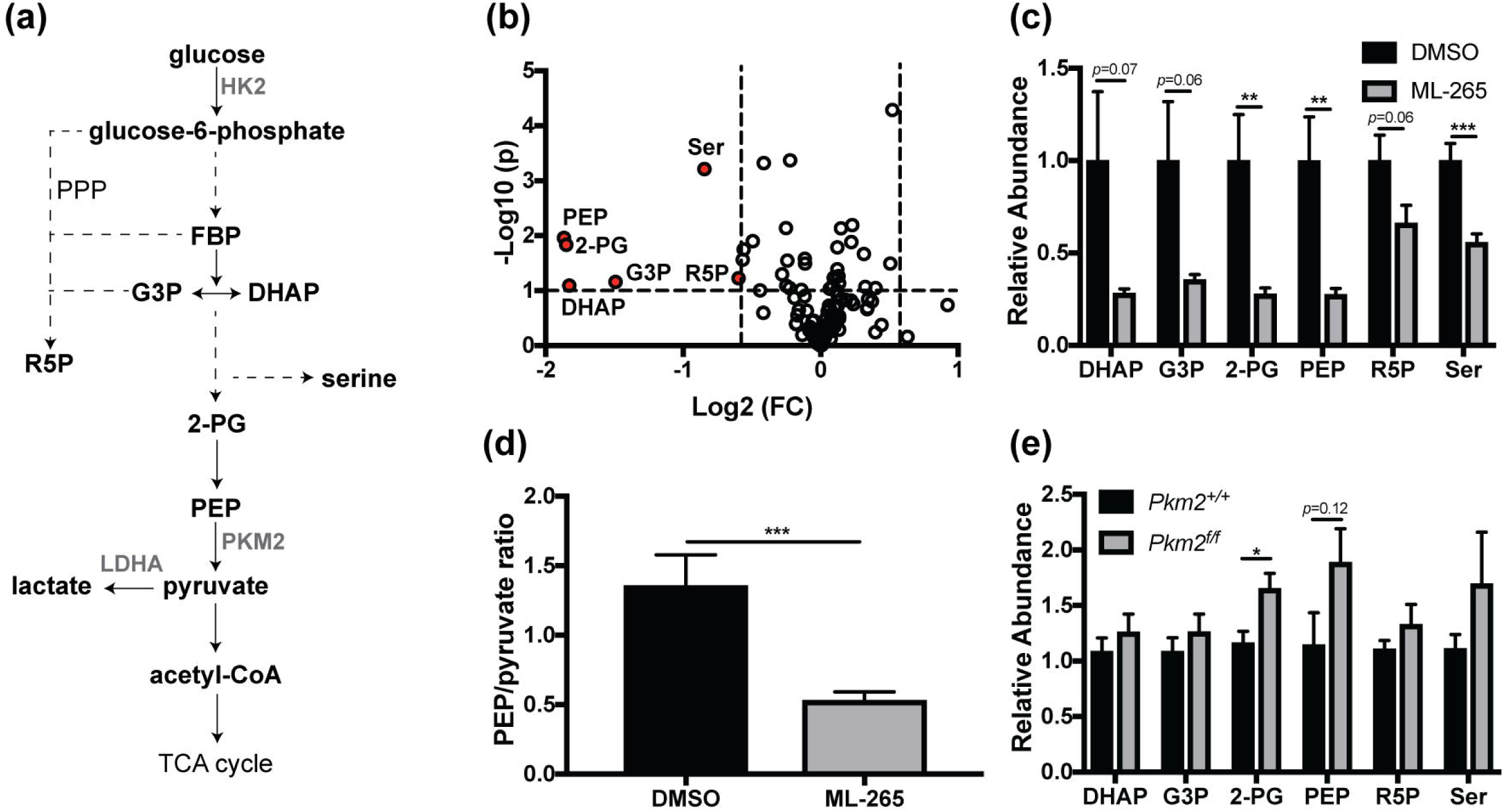
ML-265 treatment and genetic PKM1 isoform switching results in different metabolic alterations. (a) Glycolytic pathway and its interconnection with ribose 5-phosphate and serine biosynthesis. (b) 661W cells were treated with DMSO or 2 μM ML-265 for 36 hours and normalized signal intensities were determined by targeted LC-MS/MS. Volcano plot showing significantly altered metabolites after ML-265 treatment (p<0.1 and fold change >1.5 were considered significant). The – log_10_ of p value was plotted against the log_2_ of fold change (FC). Down-regulated metabolites in ML-265-treated cells compared to DMSO-treated cells are depicted in red. (c) Significantly altered metabolites in 661W cells after ML-265 treatment. Relative abundance is the ion intensity relative to DMSO-treated cells. (d) Intracellular ratio of phosphoenolpyruvate (PEP) to pyruvate, which are a substrate and a product, respectively, of the PKM2-catalyzed reaction. n=12 biological replicates per treatment. (e) Relative abundance of glycolytic and biosynthetic intermediates in the retinas of the rod, photoreceptor-specific, *Pkm2* conditional knockout mice (Rho-Cre:*Pkm2*^*f/f*^) versus retinas where both *Pkm2* alleles are present (Rho-Cre:*Pkm2*^+*/*+^). Relative abundance is the ion intensity relative to that observed in the *Pkm2*^+*/*+^ mice. n=4-6 animals per group. Mean ± SEM; *, p<0.05; **, p<0.01; ***, p<0.005. PPP, pentose phosphate pathway; PEP, phosphoenolpyruvate; 2-PG, 2-phosphoglycerate; G3P, glyceraldehyde 3-phosphate; DHAP, dihydroxyacetone phosphate; R5P, ribose 5-phosphate; Ser, serine; TCA, tricarboxylic acid; HK2, hexokinase 2; LDHA, lactate dehydrogenase A.

### ML-265 treatment circumvents apoptosis and prevents cell death *in vitro*

Next, we wanted to investigate the role of pharmacologic activation of PKM2 and the associated metabolic alterations as a novel photoreceptor neuroprotection strategy using a disease-relevant, *in vitro* model. Previous studies have shown that Fas signaling plays a crucial role in caspase activation and photoreceptor apoptosis *in vivo* after retinal detachment.^32,33^661W cells undergo caspase-mediated cell death,^25,27,34^ and treatment of 661W cells with Fas-ligand (FasL) has been shown to lead to caspase activation and cell death similar to that observed after experimental retinal detachment, making this a good *in vitro* model of outer retinal stress.^15,35^ As such, 661W cells were treated with FasL and either ML-265 or vehicle (DMSO). When treated with ML-265 alone, no statistically significant effect on caspase activation or cell viability was found as compared to vehicle (DMSO) (Fig. 7a-c). When treated with FasL alone, a statistically significantly increase in both caspase 8 and 3&7 activation was observed as well as a decrease in cell viability, as has been shown previously (Fig. 7a-c).^35,36^ Treatment with ML-265 significantly decreased caspase 8 and 3&7 activation in the presence of FasL (Fig. 7a and b). In agreement with this effect on caspase activation, ML-265 treatment of 661W cells significantly increased cell viability as compared to vehicle in the presence of FasL (Fig. 7c). These results demonstrate that ML-265 treatment can circumvent the Fas proapoptotic pathway in this disease-relevant, *in vitro* photoreceptor model.

**Figure 7.**
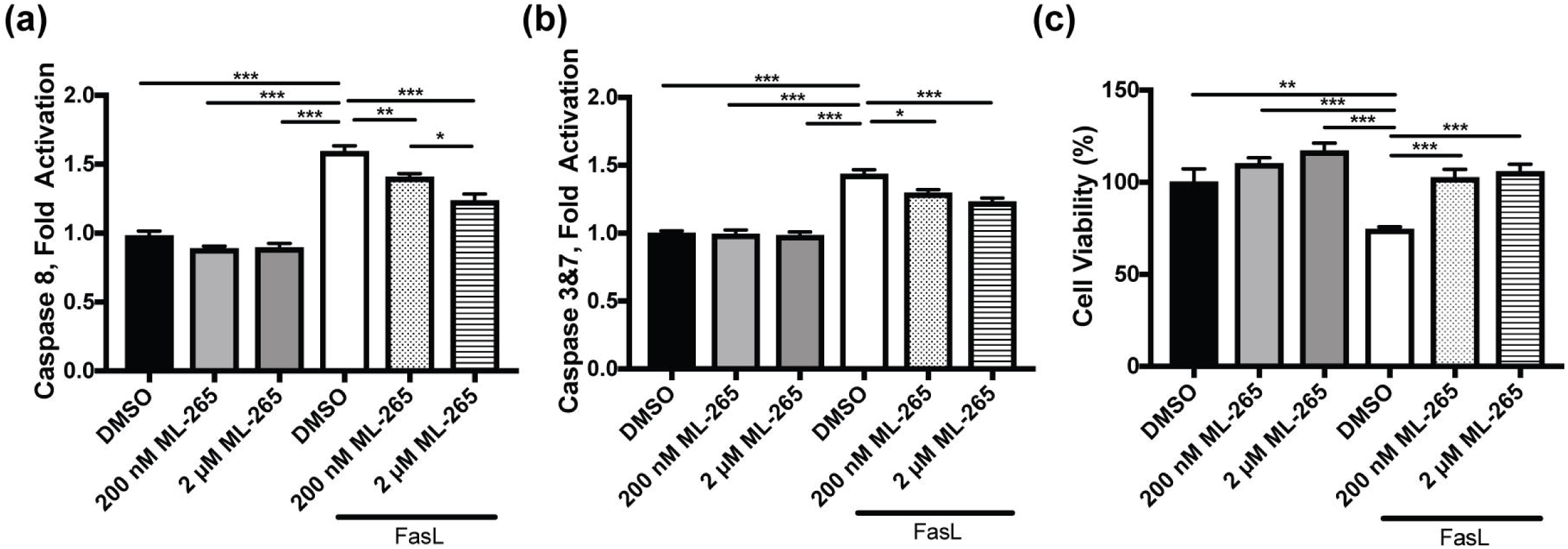
ML-265 treatment blocks caspase activation and improves cell survival in a model of apoptosis *in vitro*. Caspase 8 activity (a), caspase 3/7 activity (b), and cell viability (c) assayed in the presence of DMSO or different concentrations of ML-265 under standard tissue culture conditions or 500ng/mL FasL and 250 ng/mL hemagglutinin. Cells were pre-treated with ML-265 or DMSO for 2 hours prior to treatment with FasL, and caspase activity and cell viability were measured 8 and 48 hours, respectively, thereafter. n=8; Mean ± SEM; *, p<0.05; **, p<0.01; ***, p<0.005.

### ML-265 treatment decreases apoptosis *in vivo*

We have previously shown that Rho-Cre:*Pkm2*^*f/f*^ mice, which have a PKM2 to PKM1 isoform switch in the rod photoreceptors, demonstrate reduced entrance into the apoptosis cascade and improved photoreceptor survival in an experimental model of retinal detachment.^15^ Additionally, the *in vitro* data above suggests that ML-265 is capable of reducing Fas-mediated caspase activation. Therefore, we hypothesized that ML-265-mediated activation of PKM2 would circumvent apoptosis in a similar *in vivo* experimental model of retinal detachment in rats.^32,33^ Photoreceptor apoptosis in these experimental retinal detachments is associated with caspase activation, which has been shown to increase 3 days after detachment, and the caspase activation corresponds with the extent of photoreceptor cell death after experimental retinal detachment.^32,35,37^ Thus, rat retinas were detached, different concentrations of ML-265 or its vehicle, DMSO, were injected at the time of detachment, and the retinas were harvested after 3 days and assayed for caspase activation. Analogous to the *in vitro* results (Fig. 7a and b), retinal detachment-induced caspase 8 activation was inhibited by ML-265 treatment in a dose-dependent manner as compared to DMSO treated eyes (Fig. 8a). Likewise, ML-265 statistically significantly reduced caspase 3/7 activation at the two higher doses tested as compared to DMSO (Fig. 8b).

**Figure 8.**
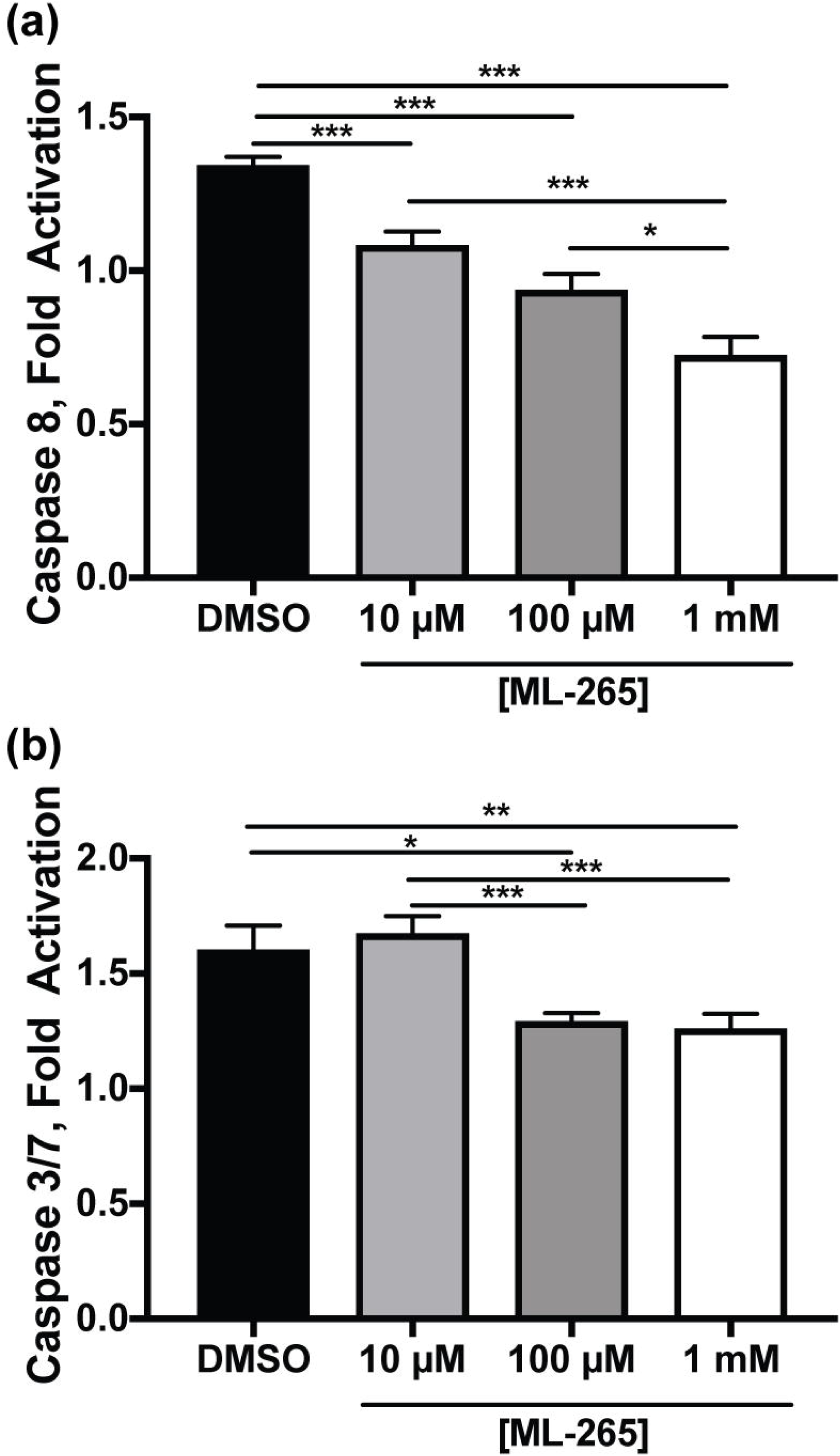
ML-265 treatment decreases caspase activation in experimental retinal detachment. Caspase 8 activity (a) and caspase 3/7 activity (b) in detached retinas after 3 days as detected by luminescent assay. Caspase activity in the detached retina was normalized to that in the attached control. ML-265 concentrations represent final vitreous concentrations after intravitreal injection. n=6 animals per group, Mean ± SEM; *, p<0.05; **, p<0.01; ***, p<0.005.

## Discussion

In this study, we show that ML-265 is capable of increasing total PK activity *in vitro* in the 661W cell culture model as well as *in vivo* in rat retinas. ML-265 has a relatively long half-life in the eye following intravitreal injection and remains active at least two weeks after injection. Extended exposure to ML-265 had no effect on photoreceptor function or survival under baseline conditions compared to vehicle-treated eyes. Interestingly, ML-265 increased PK activity without affecting the expression of genes involved in glucose metabolism. ML-265 treatment did, however, alter glucose metabolism to produce a more catabolic state. Notably, ML-265-mediated PKM2 activation reduced entrance into the apoptosis cascade in *in vitro* and *in vivo* models of outer retinal stress. Taken together, these data suggest that pharmacologic reprogramming of photoreceptor metabolism by PKM2 activators may be a novel neuroprotective strategy.

Previous studies from our laboratory and others generated and characterized a rod photoreceptor-specific, *Pkm2* conditional knockout mouse model (Rho-Cre:*Pkm2*^*f/f*^) where the loss of *Pkm2* in rod photoreceptors led to the compensatory expression of the *Pkm1* isoform.^12,15^ Rajala et al. demonstrated no significant differences in the ONL but observed that loss of PKM2 in the photoreceptors of their mice reduced the thickness of the photoreceptor end tips at 5 months of age.^12^ We observed small degenerative changes in outer retinal thickness and ONL cell count and area without statistically significant alterations in the OSEL of our Rho-Cre:*Pkm2*^*f/f*^ mice as early as 2-3 months of age.^15^ Additionally, *in vivo* electroporation of photoreceptors with a PKM1-expressing construct displayed a reduction in outer segment length within 7 weeks after birth.^10^ In contrast, continuous small molecule activation of PKM2 over the course of 6 weeks did not show any statistically significant anatomic changes in the outer retina in this study (Fig. 3). While these anatomic differences might simply be due to the fact that injections were given over the course of 6 weeks and the significant differences in the outer retinal anatomy of the Rho-Cre:*Pkm2*^*f/f*^, mice took up to 5 months to present, the *in vivo* electroporation studies with PKM1-expressing constructs would go against this postulation. Reports have shown that PKM1 and PKM2 have different cellular functions, and these different actions may not be limited to their well-described glycolytic functions.^24,38,39^ For example, some studies have suggested that PKM2 can translocate to the nucleus.^23,40,41^ Another study has shown that PKM2 has a unique effect on mitochondrial biogenesis and activation of PKM2 improves mitochondrial dysfunction.^24^ Also, PKM2 is regulated by glucose metabolites and cellular signaling processes.^18^ Therefore, compensatory increase in PKM1 expression and PK activity may not be equivalent to PKM2 activation in photoreceptors. Further studies are required to probe these different cellular functions in photoreceptors.

These Rho-Cre:*Pkm2*^*f/f*^ mice also demonstrated decreased amplitudes on scotopic electroretinography as they aged.^12,15^ After 6 weekly, intravitreal injections of small molecule PKM2 activator, no differences in scotopic or photopic electroretinography amplitudes were noted between the ML-265 and vehicle treated eyes (Fig. 4). The differences in retinal function, as assessed by electroretinography, between pharmacologic activation of PKM2 and genetic isoform reprogramming in the mouse model may again be due to the fact that injections were only given over the course of 6 weeks and significant differences in electroretinography amplitudes were not observed until after 5 months of age in the Rho-Cre:*Pkm2*^*f/f*^ mice. Yet, another possible explanation for the lack of differences between the ML-265 and vehicle treated eyes is that any small changes in retinal function in the ML-265 treated eyes may have been masked by the untoward effects of the vehicle, DMSO, on retinal function. A previous report demonstrated that DMSO displays a dose-dependent decrease in retinal function for concentrations of 0.6% or more.^42^ The low aqueous solubility of ML-265 (29.6 μg/mL) necessitates the use of organic solvents to achieve sufficient drug concentrations in the vitreous cavity due to the limiting injection volumes (2 μL).^43^ An average vitreous volume of 15 μL in rats was assumed in this study.^44,45^ Even with these low injection volumes, the retina was exposed to DMSO concentrations greater than 10% of the final vitreous volume, resulting in a decline in retinal function. Furthermore, it is unlikely that ML-265 mediated activation of PKM2 and its ability to decrease caspase activation and cell death, as shown in this study (Figs. 7 and 8), would be able to counteract the DMSO-induced toxicity. Previous studies have shown that DMSO may induce pore formation in the plasma membrane of cells and in a retinal neuronal cell line, DMSO-induced toxicity was caspase independent.^46–48^ As a result, the potential clinical applications of intravitreal ML-265 are currently limited primarily due to the low aqueous solubility and need for organic solvents. At the same time, no obvious retinal toxicity from the drug itself was observed based on the anatomical and functional analyses performed in this study. Future studies are needed to address this issue as developing novel PKM2 activators with improved solubility will be ideal for intraocular delivery.

Regardless, this study shows that pharmacologic activation of PKM2 by ML-265 increases photoreceptor resistance to apoptotic stress. Activation of PKM2 may allow the cell to optimize ATP production in the face of outer retinal stress by increasing flux through glycolysis and ultimately, shifting metabolism from an anabolic to more catabolic state as observed in this study and others.^21,24^ Notably, pharmacologic activation of PKM2 with this small molecule has also been shown to circumvent cellular apoptosis in other disease models, and it is postulated that PKM2 activation achieves this effect by reducing mitochondrial dysfunction and increasing mitochondrial biogenesis via different pathways.^24^ Furthermore, the ability of ML-265-mediated activation of PKM2 to circumvent photoreceptor apoptosis *in vivo* is in accordance with our previous results that showed decreased apoptosis and improved photoreceptor survival during acute outer retinal stress, as produced by experimental retinal detachment, in the Rho-Cre:*Pkm2*^*f/f*^ mouse model with PKM1-expressing photoreceptors.^15^ Even with this important similarity with regards to photoreceptor neuroprotection, genetic substitution of PKM1 for PKM2 and small molecule activation of PKM2 may not induce equivalent metabolic states. For example, the expression patterns of genes involved in glucose metabolism were very similar between the ML-265 and vehicle treated groups *in vitro* and *in vivo* (Fig. 5). In contrast, our knockout mouse model with selective deletion of PKM2 in photoreceptors displayed upregulation of many genes involved in glucose metabolism, including those in the non-oxidative portion of the pentose phosphate pathway and citrate shuttle.^15^ Additionally, ML-265 treatment *in vitro* resulted in decreased pools of glycolytic metabolites directly upstream of PKM2 as well as those of ribose 5-phosphate and serine, which suggests a more catabolic cellular state (Fig. 6b). Activator treatment in H1299 cells demonstrated analogous changes in the key biosynthetic intermediates, ribose 5-phosphate and serine.^21^ However, changes in these metabolite pools in the retinas of Rho-Cre:*Pkm2*^*f/f*^ mice differed from that of ML-265 treatment with 2-PG being statistically significantly increased and PEP demonstrating a similar trend (Fig. 6d). Understandably, the alterations in the metabolite pools from the Rho-Cre:*Pkm2*^*f/f*^ mice are representative of the entire retina, not just the photoreceptors, and cannot be assumed to be directly related to PK activity as we previously showed upregulation of many genes involved in central glucose metabolism in this mouse model. As a result, the differences in the observed metabolic states may reflect that PKM1 expression in photoreceptors leads to adaptive changes to compensate for the chronically elevated PK activity, unlike transient activation of PKM2 with ML-265.^21^ This data in conjunction with the understanding that PKM1 and PKM2 have different cellular functions, as discussed above, suggests that further research is needed to obtain fundamental insight into the interplay between photoreceptor metabolism and how the different PKM isoforms regulate photoreceptor survival during outer retinal metabolic stress.

In summary, we have shown that ML-265-mediated PKM2 activation and its corresponding metabolic changes reduces entrance into the apoptosis cascade in *in vitro* and *in vivo* models of acute outer retinal apoptotic stress. Studies have demonstrated that reprogramming photoreceptor metabolism may prove to be an inventive therapeutic strategy for photoreceptor neuroprotection in retinal degenerations.^4,11,49^ This study coupled with our previous work that genetically increased the levels of activated PKM in photoreceptors gives credence to this therapeutic strategy.^15^ However, further studies are needed to not only circumvent the solubility issues that limit the translation of ML-265 into the clinic but also to obtain mechanistic insight into the differential actions of PKM1 and PKM2 in photoreceptor metabolism, function, and viability.

## Materials and Methods

### Materials

#### Animals

All animals were treated in accordance with the Association for Research in Vision and Ophthalmology (ARVO) Statement for the Use of Animals in Ophthalmic and Vision Research. The protocol was approved by the University Committee on Use and Care of Animals of the University of Michigan (Protocol number: PRO00007463). Brown-Norway, adult rats were utilized for all *in vivo* ML-265 (Cayman Chemical, Ann Arbor, MI; CAS 1221186-53-3) studies except ocular pharmacokinetic studies. All rats were housed at room temperature with 12-hour light and 12-hour dark cycle. Dutch-belted rabbits were used to study the ocular pharmacokinetics of ML-265. Rod photoreceptor-specific, *Pkm2* conditional knockout mice (Rho-Cre:*Pkm2*^*f/f*^) were created by crossing mice with Lox-P sites flanking *Pkm2*-specific exon 10 (*Pkm2*^*flox/flox*^, Jackson Laboratories, Bar Harbor, ME) with Rho-Cre mice (courtesy of David Zacks, MD, PhD), in which Cre-combinase is expressed specifically in rod photoreceptors as previously described.^15,50,51^

#### Cell Culture

The 661W photoreceptor cell line was generously provided by Dr. Muayyad al-Ubaidi (Department of Cell Biology, University of Oklahoma Health Sciences Center, Oklahoma City, OK, USA).^27^ 661W cells were maintained in Dulbecco’s modified Eagle’s medium (DMEM, Thermo Fisher Scientific, Waltham, MA; Cat No. 11995065) supplemented with 10% fetal bovine serum (FBS), 90 units/mL penicillin, 0.09mg/mL streptomycin, 32 mg/L putrescine, 40 µL/L of β-mercaptoethanol, and 40 µg/L of both hydrocortisone 21-hemisuccinate and progesterone. Cells were grown at 37 °C in a humidified atmosphere of 5% CO_2_ and 95% air.

#### Chemicals

All reagents were analytical grade and purchased from Sigma (St. Louis, MO). ML-265 was purchased from Cayman Chemical (Ann Arbor, MI; CAS 1221186-53-3).

### Pyruvate Kinase Activity Enzyme Assay

#### Recombinant Enzyme

A continuous, enzyme-coupled assay, which uses lactate dehydrogenase (LDH) and measures the depletion of NADH via absorbance at 340 nm was utilized to determine the pyruvate kinase activity. For AC_50_ measurements (concentration of activator necessary to achieve half-maximal activation) with ML-265, assays were performed in 96-well format using 200 μL/well assay volume with final concentrations of 20 nM human recombinant PKM2 (Sigma, SAE0021), differing concentrations of ML-265, 0.5 mM PEP, 1 mM ADP, 0.2 mM NADH, and 8 U of lactate dehydrogenase (LDH) in an assay buffer of 50 mM Tris-HCl (pH 7.4), 100 mM KCl, and 5 mM MgCl_2_ as previously described.^15,17^ The decrease in absorbance at 340 nm was monitored using a SPECTROstar Omega plate reader (BMG LABTECH Inc., Cary, NC, USA). Initial velocities were calculated with the MARS software. Data were normalized to DMSO (dimethyl sulfoxide)-treated PKM2 activity.

#### Cell culture

For 661W cell line experiments, media was replaced prior to the start of treatment with DMSO or ML-265. Cells were incubated with DMSO or different concentrations of ML-265 for 2 hours. Cells were lysed and homogenized in RIPA Lysis and Extraction Buffer (Catalog number: 89900, Life Technologies Corporation, Grand Island, NY) with protease inhibitors (Complete-Mini, Roche Diagnostics, Indianapolis, IN) and cellular debris was removed by centrifugation at 10,000 rpm for 10 minutes. Ten microliters of the supernatant was used to assess pyruvate kinase activity, and the activity was normalized to total protein content as previously described.^22^

#### Animals

Intravitreal injections of ML-265 or vehicle (DMSO) were performed in Brown-Norway, adult rats. Rodents were anesthetized with a mix of ketamine (100 mg/mL) and xylazine (20 mg/mL), and pupils were dilated with topical phenylephrine (2.5%) and tropicamide (1%). A 25-gauge microvitreoretinal blade (Walcott Rx Products, Ocean View, NJ) was used to create a sclerotomy located 1-2 mm posterior to the limbus with care taken to avoid lens damage. A blunt, 34-gauge cannula was introduced through the sclerotomy into the vitreous cavity. Two microliters of different concentrations of ML-265 or DMSO was slowly injected into the vitreous cavity. This volume was utilized as it has been shown to produce minimal reflux, and lower volumes may not produce adequate reproducibility.^44,45^ An average vitreous volume in rats of 15 μL was assumed in order to estimate intravitreal concentrations of ML-265 after injection throughout this study.^44,45^ Left eyes were injected with ML-265, and right eyes were injected with DMSO, which served as the control that pyruvate kinase activation was based upon. Four hours after intravitreal injection, retinas from experimental rat eyes were dissected from the RPE-choroid, homogenized, and lysed in RIPA Lysis and Extraction Buffer with protease inhibitors. The assay was carried out using 4 microliters of rat retinal lysate in an enzyme buffer mixture (50 mM Tris-HCl, pH 7.4, 100 mM KCl, 5 mM MgCl_2_, 1 mM ADP, 0.5 mM PEP, and 0.2 mM NADH) and 8 units of LDH similar to what has been previously described.^15,17^ Pyruvate kinase activity was normalized to total protein in each retinal lysate.

### Quantitative Real-Time PCR

661W cells were grown to 70% confluency at 37 °C in a humidified atmosphere of 5% CO_2_ and 95% air in Dulbecco’s modified Eagle’s medium containing 25 mM glucose, 10% fetal bovine serum, 300 mg/L glutamine, 32 mg/L putrescine, 40 µL/L of β-mercaptoethanol, and 40 µg/L of both hydrocortisone 21-hemisuccinate and progesterone. The media also contained penicillin (90 units/mL) and streptomycin (0.09 mg/mL). One day prior to the start of treatment, the media was replaced with fresh media containing 5.5 mM glucose. Treatment was started by replacing the media with fresh media containing 5.5 mM glucose and DMSO or 2 µM ML-265, and the respective media was replaced with fresh media every day. Cells were harvested after 3 days of treatment.

Intravitreal injections of ML-265 or DMSO were performed in Brown-Norway, adult rats as described above. Two microliters of 7.5 mM ML-265 or equal volume of DMSO was slowly injected into the vitreous cavity. Left eyes were injected with ML-265 or DMSO. The right eye served as the control, with all steps of the procedure performed, except for introduction of the blunt cannula and injection of ML-265 or DMSO. Retinas were harvested 3- and 7-days post-injection.

Total RNA was purified from retinal tissues of rats and 661W cells as previously described.^15^ Quantitative PCR was performed using the Rat Glucose Metabolism RT^2^ Profiler™ PCR Array (Qiagen, Cat No./ID: PARN-006Z) or Mouse (661W cells) Glucose Metabolism RT^2^ Profiler™ PCR Array (Qiagen, Cat No./ID: PAMM-006Z), respectively. Reactions were undertaken using a CFX96 real time PCR system (Bio-Rad Laboratories) as previously described.^15^ The RT^2^ Profiler™ PCR Array Data Analysis Template from Qiagen was utilized for all data analyses. Genes were excluded if their relative expression level was low or not detected in all experimental groups. Data were normalized to the housekeeping genes *Actb, B2m*, and *Rplp1* in rats and *Actb, B2m*, and *Gusb* in 661W cells.

### Metabolism measurements

Metabolomics experiments were performed as previously described.^29,30,52^ 661W cells were plated in 10 cm plates in DMEM containing 25 mM glucose, 10% fetal bovine serum, 300 mg/L glutamine, 32 mg/L putrescine, 40 µL/L of β-mercaptoethanol, and 40 µg/L of both hydrocortisone 21-hemisuccinate and progesterone. Twenty-four hours prior to the initiation of treatment, the media was changed to DMEM with 5.5 mM glucose, 4 mM glutamine, 10% fetal bovine serum, and without sodium pyruvate. At t=0, the media was refreshed, and the cells were treated with DMSO or 2 μM ML-265 for 36 hours. The cells were extracted with 80% (vol/vol) methanol equilibrated at −80 degrees Celsius. Metabolite fractions were normalized to protein concentration obtained in a parallel 10 cm plate for each treatment condition. The extracts were lypholized by SpeedVac (Thermo Fisher Scientific) and analyzed via targeted liquid chromatography-tandem mass spectrometry (LC-MS/MS) with an Agilent Technologies Triple Quad 6470 instrument.

Mouse retinas from 4-month-old rod photoreceptor-specific, *Pkm2* conditional knockout mice (Rho-Cre:*Pkm2*^*f/f*^) or mice with both *Pkm2* alleles present (Rho-Cre:*Pkm2*^+*/*+^) were harvested for metabolite extraction as previously described.^15,53,54^ Briefly, retinas were harvested from each eye of the animal, combined, and snap-frozen. For metabolite measurement by LC-MS/MS, tissue was extracted in 80% methanol equilibrated at −80° Celsius, homogenized, and lypholized by SpeedVac. These samples were analyzed by targeted LC-MS/MS via dynamic multiple reaction monitoring (dMRM). The data were pre-processed with Agilent MassHunter Workstation Quantitative Analysis Software (B0900). The pre-processed data were then post-processed for further quality control. Each sample was normalized by the total intensity of all metabolites and then each metabolite abundance level was finally normalized to the median of all abundance levels for comparisons, statistical analyses, and visualizations. The statistical significance test was done by a two-tailed t-test with a significance threshold level of 0.1.^29,30^

### Caspase Activity Assay

Luminescent assay kits (Caspase-Glo 8 and 3/7 Assay Systems, Promega, Madison, WI, Cat Nos. G8200 and G8090, respectively) were utilized to measure caspase 8 and caspase 3/7 activity according to the manufacturer’s instructions. The 661W cells were seeded in white-walled 96-well plates at 2,500 cells/well for 24 hours prior to treatment. Cells were pre-treated with ML-265 or DMSO for 2 hours prior to treatment with 500 ng/ml FasL (Recombinant Mouse Fas Ligand/TNFSF6 Protein, Cat No. 6128-SA-025, R&D Systems Inc., Minneapolis, MN, USA) and 250 ng/ml HA (Hemagglutinin/HA Peptide Antibody, Cat No. MAB060, R&D Systems Inc., Minneapolis, MN, USA). Caspase activity was measured at 8 hours status post treatment by incubating the cells with substrate for 1 hour according to the manufacturer’s instructions. Luminescence was measured in a plate reader luminometer (BMG Labtech, Inc, Cary, NC).

### Cell viability

Cell viability was measured using a luminescent assay kit not based on ATP (RealTime-Glo MT Cell Viability Assay, Promega, Madison, WI, Cat No. G9711). 661W cells were seeded in 96-well plates (Nunc, Rochester, NY) at 2,500 cells/well 24 hours prior to treatment. The cells were then treated with DMSO or different concentrations of ML-265. Cell viability was measured 48 hours after treatment according to the manufacturer’s instructions. Additionally, 661W cells were pre-treated with ML-265 or DMSO for 2 hours prior to the addition of 500 ng/mL FasL and 250 ng/mL HA. Viability was again assessed at 48 hours after addition of FasL and HA. Luminescence was measured in a plate reader luminometer (BMG Labtech, Inc, Cary, NC).

### Pharmacokinetics

Pharmacokinetic studies were performed in Dutch-belted rabbits. As previously described, male rabbits were anesthetized with inhalant anesthesia (sevoflurane) prior to each aqueous humor collection or intravitreal injection. An external heat source was used to prevent hypothermia. Eyes were anesthesized with proparacaine and dilated with 1% tropicamide and 2.5% phenylephrine ophthalmic drops. A Flynn pediatric lid speculum was placed in the eye and betadine was applied to the injection site. The temporal sclera was marked with a caliper 2 mm from the limbus. A 30-gauge needle was inserted into the mid-vitreous, and 50 μL of 2.3 mM or 23 mM ML-265 was administered intravitreally. Delivery of the compound into the eye was confirmed via clinical inspection.^55^ An aqueous humor sample (100 μL) was collected from each eye prior to intravitreal injection to serve as a baseline control. A 30-gauge needle on a 1 ml syringe was used to enter the anterior chamber of the eye and aspirate the aqueous humor in a controlled fashion. Following the administration of ML-265, aqueous sample collections (100 μL) were performed at predetermined intervals, including 0, 3, 6, 24, 48, 168, and 336 hours. Samples were analyzed by LC-MS/MS.

### Toxicity

Weekly intravitreal injections of 2 μL of 7.5 mM ML-265 or 2 μL of DMSO were performed in Brown-Norway, adult rats as described above for a total of 6 weeks. Optical coherence tomography, as described below, was utilized to measure retinal thickness at baseline immediately prior to the first intravitreal injection and again, one week after the sixth and final injection (6 weeks in total after the baseline measurement). After this final *in vivo* assessment, rats were euthanized, and the eyes were enucleated for *ex vivo* analysis.

### Functional Assessment Testing

#### Optical Coherence Tomography

A spectral domain ophthalmic imaging system (SD-OCT, Bioptigen, Morrisville, NC) was used to non-lethally measure retinal thickness in adult rats. The rats received 0.5% tropicamide drops to stimulate eye dilation. The rats were anesthetized using an intraperitoneal injection of ketamine (50 mg/kg bodyweight) and xylazine (5 mg/kg bodyweight). One horizontal and one rectangular B-scan was done on each eye, of each animal. The scans were processed using the Bioptigen Diver software. Outer retinal thickness measurements (inner-most aspect of outer plexiform layer to inner aspect of retinal pigment epithelium) and outer segment equivalent length measurements (OSEL, inner segment/outer segment junction to retinal pigment epithelium inner surface) were obtained at four points, 0.35mm from the optic nerve head according to the template in the Diver software.^15,56,57^ The average of the 4 values was used for analysis.

#### Electroretinography

A Diagnosys Espion E^2^ Electrophysiology System (Diagnosys, Lowell, MA) was used to assess retinal function. The rats received a drop of 0.5% tropicamide to stimulate eye dilation and a drop of 0.5% proparacaine to numb the eyes. The rats were anesthetized using an intraperitoneal injection of ketamine (50 mg/kg bodyweight) and xylazine (5 mg/kg bodyweight). The electroretinogram (ERG) was recorded using a small contact lens that is lightly placed on the surface of the cornea; it was cushioned with a drop of 2.5% Goniosol. A scotopic protocol was conducted using a stimulus intensity of 12.31 cd*s/m^2^. A 10-minute light adaption period preceded the photopic protocol. The photopic protocol utilized a stimulus intensity of 34.19 cd*s/m^2^. During testing, body temperature was maintained at 37-38 degrees Celsius by a heating element.

### Experimental Model of Retinal Detachment

Detachments were created in Brown-Norway, adult rats as previously described.^15,32,35,37,58^ Briefly, rodents were anesthetized with a mix of ketamine (100 mg/mL) and xylazine (20 mg/mL), and pupils were dilated with topical phenylephrine (2.5%) and tropicamide (1%). A 25-gauge microvitreoretinal blade (Walcott Rx Products, Ocean View, NJ) was used to create a sclerotomy located 1-2 mm posterior to the limbus with care taken to avoid lens damage. A subretinal injector was introduced through the sclerotomy into the vitreous cavity and then through a peripheral retinotomy into the subretinal space. Sodium hyaluronate (10 mg/mL) (Abbott Medical Optics, Healon OVD) was slowly injected to detach the neurosensory retina from the underlying retinal pigment epithelium (RPE). In all experiments, approximately one-third to one-half of the neurosensory retina was detached. ML-265 or vehicle, DMSO, were injected into the subretinal space via the same peripheral retinotomy immediately after creation of the detachment as previously described.^35^ Detachments were created in the left eye. The right eye served as the control with all the steps of the procedure performed, except for introduction of the subretinal injector and injection of sodium hyaluronate.

ML-265 and DMSO-treated detached retinas were harvested after 3 days to determine caspase activity. As previously described, the tissue was homogenized using a sonicator at 20% power in buffer consisting of 20 mM MOPS, pH 7.0, 2mM EGTA, 5mM EDTA, 0.1% Triton X-100, 1 tablet of protease inhibitor (Complete Mini; Roche Diagnostics, Indianapolis, IN) and 1 tablet of phosphatase inhibitor (PhosSTOP; Roche Diagnostics, Indianapolis, IN). The homogenized samples were centrifuged at 10,000 *g* for 10 minutes at 4°C to remove cellular debris. Protein concentration was determined using a Micro BCA protein assay kit (ThermoScientific, Rockford, IL; Cat No. 23235). Substrate was incubated with 50 μg of protein from the retinal lysates per the manufacturer’s instructions and luminescence was measured. Caspase activity in the detached retina was normalized to that of the attached retina.^15^

### Histology

Rats were euthanized and the eyes were enucleated after 6 weekly intravitreal injections of ML-265 or DMSO. The eyes were fixed overnight in paraformaldehyde, embedded in paraffin, and placed in a tissue processor (Tissue-Tek II; Sakura, Tokyo, Japan). Eyes were sectioned (6 μm) using a standard paraffin microtome (Shandon AS325, Thermo Scientific, Cheshire, England). These sections were stained with hematoxylin and eosin for outer nuclear layer (ONL) cell counts and retinal area measurements.^15^

### Cell Counts and Retinal Area Measurements

A Leica DM6000 microscope (Leica Microsystems Wetzlar, Germany) was utilized for retinal images. As previously described, the total number of cells in the ONL and the total area of the retina and ONL (from the outer edge of the ONL to the inner limiting membrane) were measured using a macro program in ImageJ on entire sections through the plane of the optic nerve.^15,59^ Photoreceptor inner and outer segments were not included in the total retinal area measurement due to retraction or stretching that can occur during processing. Differences in angles of sectioning and inter-sample comparisons were accounted for by normalizing ONL cell counts or ONL area to the inner retinal area (total retinal area minus ONL area) of each section. Likewise, as ML-265 is specific for PKM2, which resides in the outer retina, any inner retinal area changes secondary to the procedure or vehicle should be consistent between both treatments examined.^12,15,17^ Data are represented as mean ± SEM.

### Data Analysis

Results are expressed as mean ± SEM. Data was analyzed using Student *t* test or one-way ANOVA followed by Bonferroni post hoc test. Prism 7.0 (GraphPad Software, San Diego, CA) was used for all statistical analysis.

### Data Availability

The datasets generated during and/or analyzed during the current study are available from the corresponding author on reasonable request.

## Acknowledgements

This study was supported by grants from the NEI 5K08EY023982-03, Research to Prevent Blindness, VitreoRetinal Surgery Foundation, and utilized the Vision Research Core funded by P30 EY007003 from the National Eye Institute. C.A.L. was supported by a Dale F. Frey Award for Breakthrough Scientists from the Damon Runyon Cancer Research Foundation (DFS-09-14); Junior Scholar Award from The V Foundation for Cancer Research (V2016-009); and a Kimmel Scholar Award from the Sidney Kimmel Foundation for Cancer Research (SKF-16-005). Metabolomics studies performed at the University of Michigan were supported by NIH grant DK097153.

## Author Contributions

T.W. and C.B. designed the research. T.W., M.P., E.W., A.S., P.S., L.Z., L.D., and H.H. performed the research. T.W., M.P., E.W., P.S., L.D., M.P., and C.L. analyzed the data. T.W. and C.B. wrote the paper with assistance from E.W., M.P., and C.L.

