## Supplementary material for "Small molecule activation of metabolic enzyme pyruvate kinase muscle isozyme 2, PKM2, provides photoreceptor neuroprotection": Table S1, Supplementary Information

**Table S1. List of metabolites and their parameters detected by LC/MS.**

| Compound Name | Precursor Ion (Da) | Product Ion (Quantifier) (Da) | Product Ion (Qualifier) (Da) | Retention Time (min) | Collision Energy (Volts) |
|---|---|---|---|---|---|
| ±-Mevalonolactone | 189.1 | 59.1 | | 2.91 | 6 |
| 2-2-Dimethyl Succinic acid | 145.1 | 101.1 | | 14 | 11 |
| | 145.1 | | 127.1 | 14 | 8 |
| 2-3-Dihydroxybenzoic acid | 153 | 109 | | 14.5 | 15 |
| | 153 | | 53.2 | 14.5 | 25 |
| 2-3-Dihydroxyisovalerate | 133.1 | 75.1 | | 10.8 | 10 |
| | 133.1 | | 57.2 | 10.8 | 15 |
| 2-3-Pyridinedicarboxylic acid | 166 | 122 | | 14.9 | 6 |
| | 166 | | 78.1 | 14.9 | 13 |
| 2-4-Quinolinediol | 160 | 117.9 | | 15.2 | 19 |
| | 160 | | 42.2 | 15.2 | 29 |
| 2-Deoxyadenosine | 250 | 134.05 | | 6.5 | 12 |
| | 310.1 | 250 | | 6.5 | 4 |
| 2-Deoxyadenosine 5-diphosphate | 410 | 79.1 | | 15.7 | 44 |
| | 410 | | 158.7 | 15.7 | 24 |
| 2-Deoxyadenosine 5-monophosphate | 330 | 79 | | 12.7 | 48 |
| | 330 | | 134.05 | 12.7 | 28 |
| 2-Deoxycytidine | 286.1 | 226 | | 2 | 4 |
| | 286.1 | | 93 | 2 | 20 |
| 2-Deoxycytidine 5-diphosphate | 386 | 79 | | 14.7 | 36 |
| | 386 | | 97 | 14.7 | 20 |
| 2-Deoxycytidine 5-monophosphate | 306 | 79 | | 10.4 | 44 |
| | 306 | | 194.9 | 10.4 | 14 |
| 2-Deoxy-D-glucose 6-phosphate | 243 | 79.1 | | 9.1 | 45 |
| | 243 | | 97 | 9.1 | 14 |
| 2-Deoxy-D-ribose | 193.1 | 59.2 | | 1.4 | 16 |
| | 193.1 | | 133 | 1.4 | 0 |
| 2-Deoxyguanosine | 266.1 | 150 | | 4 | 12 |
| | 266.1 | | 108 | 4 | 36 |
| 2-Deoxyguanosine 5-diphosphate | 426 | 79.1 | | 15.4 | 48 |
| | 426 | | 158.9 | 15.4 | 28 |
| 2-Deoxyguanosine 5-monophosphate | 346 | 79.1 | | 12.6 | 48 |
| | 346 | | 97 | 12.6 | 36 |
| 2-Deoxyinosine | 251.1 | 135 | | 3.7 | 20 |
| | 251.1 | | 108 | 3.7 | 42 |
| 2-Deoxyribose 5-phosphate | 213 | 79.1 | | 9.23 | 40 |
| | 213 | | 96.9 | 9.23 | 12 |
| 2-Deoxyuridine | 227.1 | 184 | | 2.7 | 6 |
| | 227.1 | | 42.2 | 2.7 | 20 |
| 2-Deoxyuridine 5-triphosphate | 467 | 158.8 | | 16.8 | 36 |
| | 467 | | 368.9 | 16.8 | 20 |
| 2-Isopropylmalic acid | 175.1 | 113 | | 15.5 | 13 |
| | 175.1 | | 115 | 15.5 | 13 |
| 2-Ketobutyrate | 101 | 57.2 | | 12.4 | 4 |
| 2-Methyl-1-butanol | 87.1 | 43.3 | | 9.6 | 5 |
| 2-Phosphoglyceric acid | 185 | 97 | | 14.8 | 14 |
| | 185 | | 79 | 14.8 | 37 |
| 3-2-Hydroxyethylindole | 160.1 | 130.06 | | 15.4 | 14 |
| | 160.1 | | 142.07 | 15.4 | 16 |
| 3-Dehydroshikimic acid | 171 | 127 | | 7.4 | 8 |
| | 171 | | 109 | 7.4 | 17 |

| | | | | | |
|---|---|---|---|---|---|
| 3-Hydroxyanthranilic acid | 152 | 108 | | 12.1 | 12 |
| | 152 | | 107 | 12.1 | 23 |
| 3-Hydroxy-DL-kynurenine | 223.1 | 205.9 | | 2.3 | 4 |
| | 223.1 | | 162 | 2.3 | 10 |
| 3-Hydroxyphenylacetic acid | 151 | 107.1 | | 14.2 | 4 |
| | 151 | | 65.2 | 14.2 | 26 |
| 3-Indoleacetic acid | 174.1 | 130 | | 16.5 | 7 |
| | 174.1 | | 128 | 16.5 | 19 |
| 3-Methylglutaric acid | 145.1 | 101.1 | | 14.1 | 11 |
| | 145.1 | | 41.3 | 14.1 | 52 |
| 4-Aminobenzoic acid | 136 | 92 | | 9.8 | 8 |
| 4-Guanidobutyric acid | 144.1 | 102 | | 1.4 | 8 |
| | 144.1 | | 41.2 | 1.4 | 32 |
| 4-Hydroxybenzoic acid | 137 | 93 | | 12.1 | 14 |
| | 137 | | 65.2 | 12.1 | 34 |
| 4-Hydroxy-L-glutamic acid | 162 | 144 | | 5.3 | 6 |
| | 162 | | 72.1 | 5.3 | 16 |
| 4-Hydroxyphenyl-pyruvic acid | 179 | 107 | | 14.7 | 4 |
| 4-Methyl-2-oxovaleric acid | 129.1 | 85.1 | | 13.7 | 7 |
| 4-Pyridoxic acid | 182 | 138 | | 15.3 | 12 |
| | 182 | | 108.1 | 15.3 | 20 |
| 4-Quinolinol | 144 | 66.1 | | 9.2 | 44 |
| | 144 | | 116 | 9.2 | 28 |
| 5-Deoxy-5-(methylthio)adenosine | 356.1 | 134 | | 11.8 | 12 |
| | 356.1 | | 296 | 11.8 | 4 |
| 5-Hydroxy-3-indoleacetic acid | 190.1 | 144 | | 13 | 21 |
| | 190.1 | | 116 | 13 | 46 |
| 5-Methoxytryptamine | 189.1 | 144 | | 2.4 | 28 |
| | 189.1 | | 174 | 2.4 | 12 |
| 6-Hydroxynicotinic acid | 138 | 94.1 | | 9 | 10 |
| | 138 | | 42.3 | 9 | 27 |
| Adenine | 134 | 107 | | 2.8 | 18 |
| | 134 | | 92.1 | 2.8 | 20 |
| Adenosine | 266 | 134 | | 6.4 | 10 |
| | 326.1 | 134 | | 6.4 | 20 |
| Adenosine 3-5-cyclic monophosphate | 328 | 134 | | 13.4 | 24 |
| | 328 | | 79 | 13.4 | 48 |
| Adenosine 5-diphosphate | 426 | 159 | | 15.4 | 28 |
| | 426 | | 328 | 15.4 | 16 |
| Adenosine 5-monophosphate | 346 | 79 | | 11.6 | 38 |
| | 346 | | 97 | 11.6 | 24 |
| Adenosine 5-triphosphate | 506 | 408.1 | | 17.2 | 22 |
| | 506 | | 159 | 17.2 | 38 |
| Adenylosuccinic acid | 462 | 97 | | 16.4 | 24 |
| | 462 | | 134.05 | 16.4 | 48 |
| Adipic acid | 145.1 | 101.1 | | 13.9 | 11 |
| | 145.1 | | 83.1 | 13.9 | 12 |
| AICAR | 257.1 | 125 | | 2.3 | 10 |
| | 257.1 | | 42.2 | 2.3 | 44 |
| Allantoin | 157 | 42.2 | | 1.3 | 16 |
| | 157 | | 97 | 1.3 | 12 |
| alpha-D(+)Mannose 1-phosphate | 259 | 79 | | 7.6 | 48 |
| | 259 | | 97 | 7.6 | 14 |
| alpha-D-Glucose-1-phosphate | 259 | 79 | | 7.6 | 28 |
| | 259 | | 240.9 | 7.6 | 9 |
| alpha-Ketoglutaric acid | 145 | 101 | | 14.3 | 5 |

| | | | | | |
|---|---|---|---|---|---|
| | 145 | | 57.2 | 14.3 | 9 |
| Arabinose-5-phosphate | 229 | 79 | | 8.4 | 36 |
| | 229 | | 97 | 8.4 | 8 |
| Argininosuccinic acid | 289 | 88 | | 4.6 | 36 |
| | 289 | | 131.1 | 4.6 | 27 |
| beta-Nicotinamide adenine dinucleotide | 662.1 | 540 | | 9.5 | 12 |
| | 662.1 | | 327.9 | 9.5 | 36 |
| beta-Nicotinamide mononucleotide | 333 | 251.1 | | 3.7 | 12 |
| | 333 | | 135.1 | 3.7 | 36 |
| Cellobiose | 341.1 | 161 | | 1.3 | 5 |
| Chorismic acid | 225 | 189 | | 6.7 | 6 |
| cis-Aconitic acid | 173 | 129 | | 15.5 | 4 |
| | 173 | | 85.1 | 15.5 | 8 |
| Citramalic acid | 147 | 85.1 | | 13.8 | 12 |
| | 147 | | 87 | 13.8 | 14 |
| Citric acid | 191 | 111.01 | | 14.9 | 10 |
| | 191 | | 87.01 | 14.9 | 16 |
| CoA | 766.1 | 408.1 | | 17.8 | 40 |
| | 766.1 | | 686.2 | 17.8 | 40 |
| Creatine | 130.1 | 88.1 | | 1.3 | 8 |
| | 130.1 | | 41.2 | 1.3 | 36 |
| Creatine phosphate | 210 | 79 | | 7 | 16 |
| | 210 | | 96.9 | 7 | 4 |
| Creatinine | 112 | 41.2 | | 1.35 | 24 |
| | 112 | | 68.1 | 1.35 | 16 |
| Cysteine | 120 | 33.2 | | 1.31 | 10 |
| Cytidine | 242.1 | 109 | | 1.8 | 8 |
| | 242.1 | | 42.2 | 1.8 | 16 |
| Cytidine 5-diphosphate | 402.01 | 79 | | 14.4 | 48 |
| | 402.01 | | 158.92 | 14.4 | 24 |
| Cytidine 5-triphosphate | 482 | 158.8 | | 17 | 40 |
| | 482 | | 79 | 17 | 40 |
| Cytidine-5-monophosphate | 322 | 79 | | 9.7 | 44 |
| | 322 | | 97 | 9.7 | 22 |
| Cytosine | 110 | 67.1 | | 1.4 | 8 |
| | 110 | | 42.2 | 1.4 | 12 |
| D-+-Galactosamine | 238.1 | 159.9 | | 1.1 | 8 |
| | 238.1 | | 100 | 1.1 | 6 |
| Deoxyadenosine 5-triphosphate | 490 | 391.9 | | 17.2 | 24 |
| | 490 | | 158.9 | 17.2 | 32 |
| Deoxycytidine 5-triphosphate | 466 | 158.9 | | 16.8 | 28 |
| | 466 | | 367.9 | 16.8 | 20 |
| Deoxyguanosine 5-triphosphate | 506 | 407.9 | | 17.2 | 20 |
| | 506 | | 158.8 | 17.2 | 32 |
| Deoxythymidine 5-triphosphate | 481 | 158.8 | | 17.2 | 36 |
| | 481 | | 383.1 | 17.2 | 20 |
| D-erythro-Dihydrosphingosine | 300.3 | 199 | | 8.9 | 8 |
| | 300.3 | | 282 | 8.9 | 6 |
| D-Fructose 1,6-biphosphate | 338.9 | 97 | | 14.8 | 22 |
| | 338.9 | | 241.01 | 14.8 | 12 |
| D-Fructose 6-phosphate | 259 | 79 | | 7.6 | 48 |
| | 259 | | 97 | 7.6 | 14 |
| D-Gluconic acid | 195.1 | 75.2 | | 5.9 | 18 |
| | 195.1 | | 129 | 5.9 | 10 |
| D-Glucosamine 6-phosphate | 258 | 79 | | 1.6 | 48 |
| | 258 | | 97 | 1.6 | 16 |

| | | | | | |
|---|---|---|---|---|---|
| D-Glucose 6-phosphate | 259 | 79 | | 7.6 | 48 |
| | 259 | | 97 | 7.6 | 14 |
| Dihydroxyacetone phosphate | 169 | 79 | | 10.5 | 32 |
| | 169 | | 97 | 10.5 | 6 |
| DL-2-Aminoadipic acid | 160.1 | 142 | | 4.3 | 9 |
| | 160.1 | | 116 | 4.3 | 12 |
| DL-Glyceraldehyde 3-phosphate | 169 | 79 | | 11.7 | 28 |
| | 169 | | 97 | 11.7 | 4 |
| DL-Isocitric acid | 191 | 111.02 | | 15.1 | 11 |
| | 191 | | 173 | 15.1 | 6 |
| D-Maltose | 341.1 | 161 | | 1.3 | 5 |
| | 341.1 | | 179 | 1.3 | 4 |
| D-Mannose | 179.1 | 89 | | 1.3 | 4 |
| | 179.1 | | 59.2 | 1.3 | 16 |
| D-pantothenic acid | 218 | 88 | | 12.1 | 10 |
| | 218 | | 146.08 | 12.1 | 14 |
| D-Ribose 5-phosphate | 229 | 79 | | 8.5 | 48 |
| | 229 | | 97 | 8.5 | 10 |
| D-Ribulose 1,5-biphosphate | 308.9 | 79 | | 14.7 | 46 |
| | 308.9 | | 97 | 14.7 | 22 |
| D-Sedoheptulose-7-phosphate | 289 | 96.9 | | 8.2 | 18 |
| | 289 | | 79 | 8.2 | 48 |
| D-Xylose | 149.05 | 89.02 | | 1.2 | 4 |
| | 149.05 | | 71.01 | 1.2 | 4 |
| D-Xylulose-5-phosphate | 229 | 79 | | 8.5 | 36 |
| | 229 | | 138.98 | 8.5 | 8 |
| Epicatechin | 289.1 | 245 | | 12.2 | 11 |
| | 289.1 | | 109 | 12.2 | 26 |
| Flavin adenine dinucleotide | 784.1 | 346 | | 17 | 35 |
| | 784.1 | | 437 | 17 | 30 |
| Folinic acid | 472.2 | 315 | | 14.8 | 28 |
| | 472.2 | | 343 | 14.8 | 20 |
| Galactonic acid | 195.1 | 75.1 | | 5.9 | 18 |
| | 195.1 | | 129 | 5.9 | 11 |
| gamma-Aminobutyric acid | 102 | 84.1 | | 1.2 | 8 |
| gamma-Glu-Cys | 249.1 | 128 | | 8.1 | 8 |
| | 249.1 | | 171 | 8.1 | 9 |
| Glucoheptonic acid | 225.1 | 129 | | 5.8 | 11 |
| | 225.1 | | 75.1 | 5.8 | 18 |
| Glutathione Reduced | 306.1 | 143 | | 8.43 | 17 |
| | 306.1 | | 128.1 | 8.43 | 17 |
| Glyceric acid | 105 | 75.1 | | 5.9 | 8 |
| | 105 | | 57.2 | 5.9 | 14 |
| Glyoxylic acid | 73 | 45.2 | | 5.9 | 6 |
| Guanine | 150 | 133.02 | | 2 | 12 |
| | 150 | | 66.01 | 2 | 36 |
| Guanosine | 282.1 | 150 | | 3.9 | 17 |
| | 282.1 | | 132.9 | 3.9 | 32 |
| Guanosine 3,5-cyclic monophosphate | 344 | 150.042 | | 12.2 | 24 |
| | 344 | | 133.02 | 12.2 | 44 |
| Guanosine 5-diphosphate | 442 | 150 | | 14.8 | 31 |
| | 442 | | 158.9 | 14.8 | 20 |
| Guanosine 5-triphosphate | 522.3 | 158.9 | | 17 | 36 |
| | 522.3 | | 424 | 17 | 20 |
| Homocitrate | 205 | 125 | | 15.4 | 10 |
| | 205 | | 145 | 15.4 | 10 |

| | | | | | |
|---|---|---|---|---|---|
| Hypoxanthine | 135 | 92 | | 2 | 17 |
| | 135 | | 65.1 | 2 | 28 |
| Indoline-2-carboxylate | 162.1 | 116 | | 16.1 | 16 |
| | 162.1 | | 118 | 16.1 | 11 |
| Inosine | 267.1 | 135 | | 3.3 | 22 |
| | 267.1 | | 108 | 3.3 | 44 |
| Inosine 5-diphosphate | 427 | 158.9 | | 14.5 | 28 |
| | 427 | | 135 | 14.5 | 24 |
| Inosine 5-monophosphate | 347 | 79 | | 11.1 | 44 |
| | 347 | | 97 | 11.1 | 22 |
| Inosine 5-triphosphate | 507 | 409 | | 17.2 | 19 |
| | 507 | | 159 | 17.2 | 40 |
| Isopentenyl pyrophosphate | 245 | 209 | | 12.8 | 4 |
| Isopentyl acetate | 129.1 | 85.2 | | 14.6 | 6 |
| | 129.1 | | 41.2 | 14.6 | 12 |
| Itaconic acid | 129 | 85.1 | | 13.6 | 6 |
| | 129 | | 41.3 | 13.6 | 12 |
| Ketoisovaleric acid | 115 | 71.2 | | 16.56 | 4 |
| Ketovaleric acid | 115 | 71.2 | | 14.5 | 4 |
| Lactic acid | 89 | 43.3 | | 7.4 | 10 |
| | 89 | | 45.3 | 7.4 | 9 |
| L-Arabinose | 149 | 89.1 | | 1.4 | 4 |
| | 149 | | 59.2 | 1.4 | 12 |
| L-Arabitol | 151 | 89 | | 1.3 | 12 |
| | 151 | | 71.02 | 1.3 | 16 |
| L-Arginine | 173.1 | 131 | | 1.1 | 11 |
| | 173.1 | | 156 | 1.1 | 8 |
| L-asparagine | 131 | 113 | | 1.2 | 4 |
| | 131 | | 70 | 1.2 | 12 |
| L-Aspartic Acid | 132 | 88 | | 4.9 | 10 |
| | 132 | | 71 | 4.9 | 14 |
| L-Canavanine | 175.1 | 74.04 | | 1.1 | 10 |
| | 175.1 | | 118 | 1.1 | 8 |
| L-Carnitine | 220.1 | 146 | | 1.2 | 4 |
| L-Citrulline | 174.1 | 131 | | 1.3 | 8 |
| | 174.1 | | 42.2 | 1.3 | 48 |
| L-Cystathionine | 221.1 | 134 | | 1.3 | 8 |
| | 221.1 | | 120 | 1.3 | 9 |
| L-Cystine | 239 | 74.1 | | 1.2 | 20 |
| | 239 | | 120 | 1.2 | 4 |
| L-Dihydroorotic acid | 157 | 113 | | 7.3 | 5 |
| | 157 | | 42.2 | 7.3 | 30 |
| L-Glutamic acid | 146 | 102 | | 4.5 | 11 |
| | 146 | | 128 | 4.5 | 8 |
| L-Glutamine | 145.1 | 127 | | 1.3 | 7 |
| | 145.1 | | 109 | 1.3 | 10 |
| L-Glutathione (oxidized) | 611.1 | 306.07 | | 13.6 | 24 |
| | 611.1 | | 272.09 | 13.6 | 28 |
| L-Histidine | 154.1 | 137 | | 1.1 | 12 |
| | 154.1 | | 93.1 | 1.1 | 16 |
| L-Homocysteine | 134 | 116.9 | | 1.4 | 8 |
| L-Homocystine | 267 | 132 | | 1.5 | 8 |
| | 267 | | 115 | 1.5 | 16 |
| L-Homoserine | 118 | 72.1 | | 1.3 | 10 |
| | 118 | | 55 | 1.3 | 16 |
| L-Hydroxyglutaric acid | 147 | 128.9 | | 13.8 | 7 |

| | | | | | |
|---|---|---|---|---|---|
| | 147 | | 85.1 | 13.8 | 13 |
| Lipoamide | 204.1 | 158 | | 16.8 | 4 |
| | 204.1 | | 64.1 | 16.8 | 20 |
| L-Isoleucine | 130 | 45 | | 2.1 | 16 |
| L-Kynurenine | 207.1 | 190 | | 3.9 | 4 |
| | 207.1 | | 144 | 3.9 | 20 |
| L-Leucine | 130 | 45 | | 2.1 | 16 |
| L-Malic acid | 133 | 115 | | 13.8 | 8 |
| | 133 | | 71.1 | 13.8 | 14 |
| L-Methionine | 148 | 47.2 | | 1.8 | 12 |
| | 148 | | 100 | 1.8 | 6 |
| L-Phenylalanine | 164.1 | 147 | | 4.5 | 9 |
| | 164.1 | | 103 | 4.5 | 15 |
| L-Proline | 114 | 68.1 | | 1.3 | 12 |
| | 114 | | 45.2 | 1.3 | 12 |
| L-Serine | 104 | 74.1 | | 1.2 | 8 |
| L-Sorbose | 179.1 | 89 | | 1.3 | 4 |
| | 179.1 | | 71.02 | 1.3 | 14 |
| L-Threonine | 118 | 74.1 | | 1.3 | 9 |
| L-Tryptophan | 203.1 | 116 | | 7.9 | 14 |
| | 203.1 | | 74.1 | 7.9 | 14 |
| L-Tyrosine | 180 | 119 | | 2.3 | 15 |
| | 180 | | 93 | 2.3 | 12 |
| Maleic acid | 115 | 71.2 | | 12.8 | 7 |
| | 115 | | 27.3 | 12.8 | 15 |
| Malonic acid | 103 | 59.2 | | 12.8 | 7 |
| | 103 | | 41.3 | 12.8 | 32 |
| Melibiose | 341.1 | 179 | | 1.3 | 4 |
| | 341.1 | | 89 | 1.3 | 13 |
| Mevalonic acid | 147.1 | 128.9 | | 9.8 | 8 |
| | 147.1 | | 103.1 | 9.8 | 11 |
| Mevalonic acid 5-phosphate | 227 | 79 | | 14.8 | 40 |
| | 227 | | 97 | 14.8 | 12 |
| m-Hydroxybenzoic acid | 137 | 93.1 | | 13.4 | 10 |
| | 137 | | 65.2 | 13.4 | 28 |
| myo-Inositol | 179 | 71.02 | | 1.3 | 12 |
| | 239.1 | 179 | | 1.3 | 0 |
| N-Acetyl D-galactosamine | 220.1 | 119 | | 1.2 | 0 |
| N-Acetyl-alpha-D-glucosamine 1-phospha | 300 | 97 | | 8.7 | 16 |
| | 300 | | 79 | 8.7 | 36 |
| N-acetylaspartate | 174 | 88.1 | | 14 | 14 |
| | 174 | | 58.1 | 14 | 22 |
| N-acetylaspartylglutamate | 303.1 | 285 | | 16.1 | 6 |
| | 303.1 | | 128 | 16.1 | 14 |
| N-Acetyl-D-glucosamine 6-phosphate | 300 | 97 | | 8.8 | 18 |
| | 300 | | 79 | 8.8 | 48 |
| N-Acetylglutamic acid | 188.1 | 128.04 | | 14.1 | 10 |
| | 188.1 | | 102 | 14.1 | 16 |
| N-Acetylneuraminic acid | 308.1 | 87 | | 6.2 | 14 |
| | 308.1 | | 169.9 | 6.2 | 10 |
| N-Carbamoyl-DL-aspartic acid | 175 | 132 | | 13.5 | 7 |
| | 175 | | 88.1 | 13.5 | 19 |
| N-Carbamyl-L-glutamic acid | 189.1 | 146 | | 13.6 | 7 |
| | 189.1 | | 102.1 | 13.6 | 20 |
| N-Formyl-L-Tyrosine | 208.1 | 164 | | 12.8 | 7 |
| | 208.1 | | 107 | 12.8 | 12 |

| | | | | | |
|---|---|---|---|---|---|
| Nicotinic acid | 122 | 78.1 | | 11.9 | 11 |
| | 122 | | 51.2 | 11.9 | 32 |
| Nicotinic acid mononucleotide | 334 | 289.9 | | 8.6 | 4 |
| | 334 | | 79 | 8.6 | 40 |
| o-Hydroxy hippuric acid | 194 | 150 | | 16.5 | 11 |
| | 194 | | 121 | 16.5 | 22 |
| o-Phospho-L-Serine | 184 | 79 | | 10.2 | 36 |
| | 184 | | 97 | 10.2 | 13 |
| O-Phosphorylethanolamine | 140 | 79 | | 1.7 | 14 |
| | 140 | | 63 | 1.7 | 38 |
| Orotic acid | 155 | 42 | | 9.1 | 26 |
| | 155 | | 40 | 9.1 | 44 |
| O-Succinyl-L-homoserine | 218.1 | 117 | | 6.5 | 7 |
| | 218.1 | | 118 | 6.5 | 0 |
| Oxamic acid | 88 | 44.3 | | 6.5 | 6 |
| | 88 | | 42.3 | 6.5 | 12 |
| Phenylpyruvic acid | 163 | 91.1 | | 17.2 | 6 |
| Phosphoenolpyruvic acid | 167 | 79 | | 15.3 | 12 |
| Prephenic acid | 225 | 101 | | 12.4 | 9 |
| Pyridoxal 5 phosphate | 246 | 79 | | 14.8 | 30 |
| Pyridoxal hydrochloride | 166.1 | 138 | | 2.4 | 12 |
| | 166.1 | | 108 | 2.4 | 18 |
| Pyridoxamine | 167.1 | 121 | | 1.1 | 22 |
| | 167.1 | | 122 | 1.1 | 18 |
| Pyridoxine | 168.1 | 122.1 | | 2.4 | 17 |
| | 168.1 | | 166 | 2.4 | 9 |
| Pyruvic acid | 87 | 43.2 | | 9.5 | 4 |
| Quinic acid | 191.1 | 85.1 | | 6 | 25 |
| | 191.1 | | 93.1 | 6 | 24 |
| Riboflavin | 375.1 | 255 | | 12.6 | 14 |
| | 375.1 | | 212 | 12.6 | 28 |
| Ribonic acid gamma lactone | 147 | 59.2 | | 1.5 | 16 |
| | 147 | | 99 | 1.5 | 8 |
| S-2-Aminoethyl-L-cysteine | 163.1 | 76.1 | | 1.1 | 10 |
| | 163.1 | | 33.2 | 1.1 | 28 |
| S-5-Adenosyl-L-homocysteine | 383.1 | 134 | | 5.3 | 24 |
| | 383.1 | | 248 | 5.3 | 13 |
| Salicylic acid | 137 | 93.1 | | 16.8 | 17 |
| | 137 | | 65.2 | 16.8 | 33 |
| Shikimic acid | 173 | 111 | | 5.8 | 7 |
| | 173 | | 93.1 | 5.8 | 16 |
| Succinic acid | 117 | 73.1 | | 13 | 10 |
| | 117 | | 99 | 13 | 8 |
| Succinic semialdehyde | 101 | 57.2 | | 7.1 | 7 |
| Taurine | 124 | 80 | | 1.3 | 24 |
| Taurocholic acid | 514 | 514 | | 20.7 | 50 |
| Thiamine | 264.1 | 234 | | 1.1 | 6 |
| | 264.1 | | 148 | 1.1 | 18 |
| Thymidine | 241.1 | 42.2 | | 5.5 | 16 |
| | 241.1 | | 151 | 5.5 | 6 |
| Thymidine 5-diphosphate | 401.01 | 97 | | 15.4 | 25 |
| | 401.01 | | 79 | 15.4 | 44 |
| Thymine | 125 | 42.2 | | 2.7 | 14 |
| trans-4-Hydroxy-L-proline | 130.1 | 128 | | 1.3 | 12 |
| | 130.1 | | 66.03 | 1.3 | 16 |
| trans-Aconitic acid | 173 | 129 | | 15.55 | 4 |

| | | | | | |
|---|---|---|---|---|---|
| | 173 | | 85.1 | 15.55 | 10 |
| trans-trans Muconic acid | 141 | 53.3 | | 14.7 | 8 |
| Trehalose | 341 | 89.01 | | 1.3 | 12 |
| | 401.1 | 341 | | 1.3 | 10 |
| Trehalose 6-phosphate | 421.1 | 241 | | 7.9 | 27 |
| Uracil | 111 | 42.2 | | 1.7 | 14 |
| Uric acid | 167 | 124 | | 5.5 | 13 |
| | 167 | | 42.3 | 5.5 | 26 |
| Uridine | 243.1 | 200 | | 2.3 | 6 |
| | 243.1 | | 110 | 2.3 | 12 |
| Uridine 5-diphosphate | 403 | 158.9 | | 14.8 | 28 |
| | 403 | | 79 | 14.8 | 48 |
| Uridine 5'-diphosphogalactose | 565 | 322.9 | | 14.3 | 24 |
| | 565 | | 384.9 | 14.3 | 29 |
| Uridine 5-diphosphoglucose | 565 | 322.9 | | 14.3 | 24 |
| | 565 | | 384.9 | 14.3 | 28 |
| Uridine 5-monophosphate | 323 | 79.1 | | 10.8 | 48 |
| | 323 | | 97 | 10.8 | 24 |
| Uridine 5-triphosphate | 483 | 158.9 | | 17 | 36 |
| | 483 | | 385 | 17 | 20 |
| Vanillic acid | 167 | 123 | | 12.6 | 9 |
| | 167 | | 108 | 12.6 | 18 |
| Xanthine | 151 | 108 | | 2.1 | 16 |
| | 151 | | 42.2 | 2.1 | 24 |
| Xanthosine | 283.1 | 151 | | 8.2 | 18 |
| | 283.1 | | 108 | 8.2 | 40 |
| Xylitol | 151.1 | 89 | | 1.3 | 8 |
| | 151.1 | | 71 | 1.3 | 8 |
